# Glyco-Engineering Cell Surfaces by Exo-Enzymatic Installation of GlcNAz and LacNAz Motifs

**DOI:** 10.1101/2023.08.28.554597

**Authors:** Fabiola V. De León González, Marie E. Boddington, Martha I. Prindl, Chantelle J. Capicciotti

**Author notes:** Correspondence to: Chernoff Hall, Rm. 405 90 Bader Lane K7L 2S8 Kingston, ON, Canada. These authors contributed equally to this work (Fabiola V. De León González, Marie E. Boddington).

## Abstract

Exo-enzymatic glyco-engineering of cell-surface glycoconjugates enables the selective display of well-defined glyco-motifs bearing bioorthogonal functional groups which can be used to study glycans and their interactions with glycan-binding proteins. While the installation of monosaccharides and their derivatives using glycosyltransferase enzymes has rapidly evolved, similar strategies to introduce chemical-reporter functionalized Type 2 LacNAc motifs have not been reported. Herein, we report the chemo-enzymatic synthesis of unnatural UDP-GlcNAc and UDP-GalNAc nucleotide-sugars, and the donor and acceptor substrate tolerance of the human glycosyltransferases B3GNT2 and B4GalT1, respectively, to form derivatized LacNAc moieties. We also demonstrate that B3GNT2 can be used to exo-enzymatically install GlcNAc and GlcNAz onto cell-surface glycans. GlcNAc- or GlcNAz-engineered cells can be further extended by B4GalT1, producing LacNAc or LacNAz-engineered cells. Our glyco-engineering labeling strategy is amenable to different cell types and our work expands the exo-enzymatic glycan editing toolbox to selectively introduce unnatural Type 2 LacNAc motifs.

## Article

Cells are coated with a dense layer of glycans conjugated to proteins, lipids and small RNAs.^1^ Through interactions with glycan-binding proteins (GBPs), terminal glyco-motifs displayed on cell-surface glycoconjugates serve as ligands for GBPs to mediate a variety of physiological and pathological processes including cell-cell adhesion and communication, immune responses, and host-pathogen interactions.^2, 3^ One important motif is Type 2 *N*-acetyl-lactosamine (LacNAc), which is a disaccharide composed of Gal-β1,4-GlcNAc (Gal: galactose; GlcNAc: *N*-acetyl-glucosamine). Many GBPs use LacNAcs as their binding ligands, including galectins, which recognize the β-Gal termini of LacNAcs and are involved in cell signalling, immunomodulatory processes and cancer metastasis.^4–6^ LacNAcs are widely displayed on cells as single or multiple repeating units (polyLacNAcs) and comprise the branches of hybrid- and complex-type *N*-glycans, mucin-type *O*-glycans, and glycolipids. Additionally, LacNAc chains can act as extended scaffolds to present terminal glyco-epitopes, such as sialyl LacNAc or the Lewis antigens, away from the cell membrane. The length of the polyLacNAc chain can also influence physiological function, like maintaining immune cell homeostasis.^7^ Additionally, polyLacNAcs are upregulated in malignant tissues such as colorectal cancer, gastric carcinoma, and breast cancer.^8, 9^

In recent years, there has been an expansion of chemical tools developed to study sialylation or fucosylation of cellular glycoconjugates through metabolic or exo-enzymatic glycan labeling.^10–12^ Despite the biological importance of LacNAcs, there have been limited reports on the development of analogous tools to study this specific glyco-motif in cells. LacNAcs contain GlcNAc and Gal residues, and it is challenging to control their selective formation using commonly employed bioorthogonal chemical reporter strategies like metabolic oligosaccharide engineering (MOE). Various GlcNAc derivatives bearing chemical reporters, including azides, alkynes and photoactivatable diazirine functionalities, have been used in MOE to label cellular glycans.^13–15^ These probes have primarily been used to study *O*-GlcNAc-modified proteins since the modified UDP-GlcNAc donors biosynthesized through MOE are accepted by the nucleocytoplasmic *O*-GlcNAc transferase (OGT).^16^ However, it is possible for GlcNAc reporters to be incorporated into *N*- and *O*-linked glycans through different GlcNAc-transferases (GlcNAcTs), or as GalNAc residues through epimerases in salvage pathways and tolerance by GalNAc-transferases.^17–19^ Thus, selective display in higher-order cell-surface glycans with GlcNAc derivatives can be complicated, resulting in non-specific glycosylation and lower labeling efficiencies.^20, 21^ Notably, while Kohler and co-workers demonstrated that a diazirine-modified GlcNAc can be metabolically incorporated into *N*-glycans primarily by the GlcNAcT MGAT2, the probe was not appreciably tolerated by other transferases involved in extended LacNAc biosynthesis.^15^ With regards to the Gal residue within the LacNAc motif, to the best of our knowledge, Gal-based probes have not yet been used in MOE for incorporation and display on cell-surface glycans.

A promising alternative strategy to MOE that facilitates the specific display of glyco-motifs on cells is exogenous (exo)-enzymatic glyco-engineering. In this approach, modified nucleotide (NT)-sugars and glycosyltransferase enzymes are used to selectively edit cell-surface glycans with probes in a motif-specific manner for visualization and characterization, as well as interrogation of GBP recognition.^12^ Selectivity arises by exploiting the specificity of the glycosyltransferase employed, enabling control over the glycosidic linkage, and often the glycan subclass, that is displaying the probe-modified monosaccharide. Prominent examples have used sialyl- and fucosyl-transferases to install azide, alkyne, diazirine, biotin and other probe-modified NT-sugars onto cellular glycans.^22, 23^

In humans, extended LacNAcs are synthesized by the alternating actions of β1,3-GlcNAc-transferase and β1,4-Gal-transferase enzymes, with the GlcNAcT B3GNT2 and the Gal-transferase (GalT) B4GalT1 predominantly synthesizing polyLacNAc structures.^24, 25^ As such, extended LacNAc moieties have been glyco-engineered on erythrocytes using the human B3GNT2 and B4GalT1 enzymes with natural UDP-GlcNAc and UDP-Gal donors to install functional influenza A virus ligands.^26^ However, to our knowledge, there have been no reports of the exo-enzymatic build-up of LacNAcs on cells using unnatural NT-sugars. Additionally, the substrate tolerance of these human transferase enzymes with chemically-modified unnatural donors and acceptors has yet to be explored. Other human GlcNAcTs, including B3GNT6 and GCNT1, have been used to selectively label cell-surface *O*-GalNAc and Core 1 *O*-glycans, respectively, with an azide-bearing GlcNAz.^27^ It has also been demonstated that the bacterial β1,3-GlcNAcT NmLgtA can exo-enzymatically introduce a C6-azido UDP-GlcNAc derivative onto cells.^28^ However, in these reports, the newly installed GlcNAz or 6-azido-GlcNAc was not converted into a LacNAc moiety. With regards to the Gal residue in the LacNAc motif, the bacterial β1,4-GalT NmLgtB has been used to exo-enzymatically install a C6-azido UDP-Gal derivative into cell-surface glycans of Lec 8 CHO cells, a cell line displaying truncated *N*-glycans with exposed terminal GlcNAc acceptors.^28^

Given the important biological functions of LacNAcs, we were interested in expanding exo-enzymatic strategies to specifically engineer cell-surface glycans with LacNAc motifs bearing bioorthogonal chemical reporters. Toward this goal, we sought to prepare a small library of unnatural NT-sugar derivatives to assess their tolerance by the human LacNAc synthesizing enzymes B3GNT2 and B4GalT1, as well as demonstrate a proof-of-concept application for exo-enzymatic cell-surface glycan editing.

A common chemo-enzymatic approach to prepare UDP-GlcNAc and UDP-GalNAc analogues is using a one-pot multi-enzyme (OPME) reaction system.^29^ These reactions use three enzymes to form the NT-sugar products from free-reducing monosaccharides: an *N*-acetyl-hexosamine 1-kinase, such as *Bifidobacterium Longum* NahK, to form glycosyl-1-*O*-phosphate intermediates; GlcNAc- or GalNAc-uridylyltransferases (or pyrophosphorylases) to form the activated NT-sugar donor; and an inorganic pyrophosphatase, such as *Pasteurella multocida* PmPpA, to drive reaction progress by hydrolyzing the pyrophosphate by-product (Figure 1A). Several uridylyltransferases have been used to prepare UDP-GlcNAc and UDP-GalNAc derivatives, including *P. multocida* GlcNAc-1-*O*-phosphate uridylyltransferase (PmGlmU)^30^ and human UDP-GalNAc pyrophosphorylase AGX1.^31, 32^ As the success of OPME reactions relies on the substrate promiscuity of these enzymes, we first sought to assess a small library of unnatural GlcNAc and GalNAc derivatives for their tolerance as substrates for the preparation of UDP-GlcNAc and UDP-GalNAc derivates using PmGlmU or AGX1 (Figure 1). Modified GlcNAc and GalNAc derivatives containing the common azide (Az), alkyne (Alk), and diazirine (DAz) bioorthogonal chemical reporters on the C2 *N*-acyl side chain of the pyranose ring were synthesized by adapting previously reported protocols (Supporting Information, Scheme S1).^33–35^ Each derivative was then subjected to an OPME reaction with NahK, PmGlmU or AGX1, and PmPpA (Figure 1A), and percentage conversions into glycosyl-1-*O*-phosphate intermediates and UDP-sugar products were assessed by ^1^H-NMR spectroscopy (Figure 1C) after overnight incubation. As expected, complete conversion into the corresponding glycosyl-1-*O*-phosphate intermediate was observed for all monosaccharide analogues, confirming the broad substrate promiscuity of NahK (Figures S1-S8).^32, 36^ In general, both PmGlmU and AGX1 tolerated both GlcNAc and GalNAc-based monosaccharides with small functional groups on the C2 *N*-acyl side chain (Figure 1C). Full conversion was observed with PmGlmU to form UDP-GlcNAz (**2**), and with AGX1 to form UDP-GalNAz (**6**). AGX1 resulted in 43% conversion into UDP-GlcNAz (**2**), and while PmGlmU was capapble of forming UDP-GalNAc (**5**), it did not tolerate GalNAz-1-*O*-phosphate as a suitable substrate.

**Figure 1.**
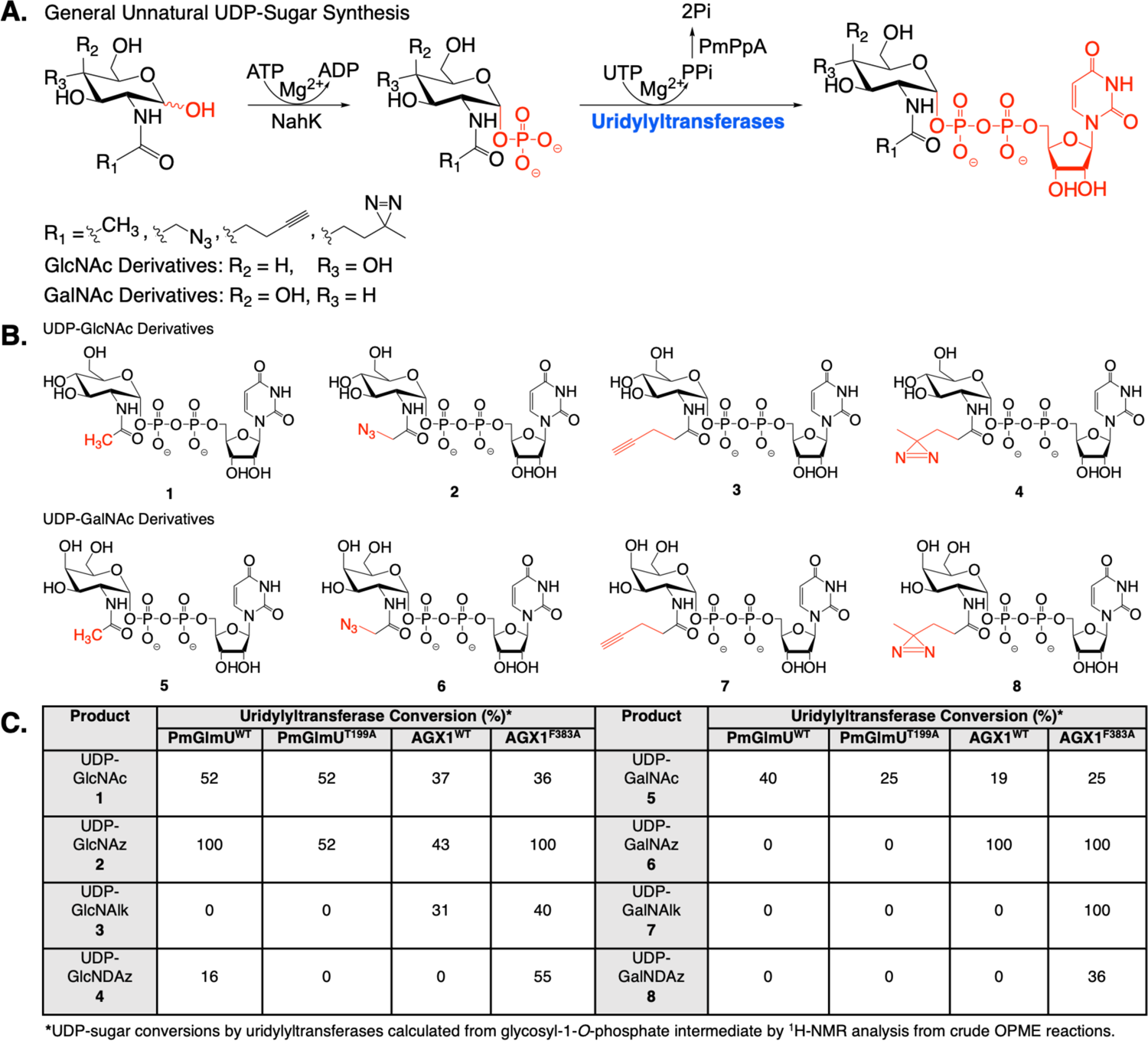
Substrate specificity of WT and Mutant PmGlmU and AGX1 uridylyltransferases. **A)** OPME synthesis of unnatural UDP-GlcNAc and UDP-GalNAc derivatives using NahK from *B. Longum*, PmPpA from *P. multocida*, and PmGlmU WT or T199A from *P. multocida*, or human AGX1 WT or F383A, enzymes. **B)** Structures of the UDP-GlcNAc (**1-4**) and UDP-GalNAc (**5-8**) derivatives synthesized using an OPME approach. **C)** Percentage conversions from glycosyl-1-*O*-phosphate intermediates to UDP-sugar products for WT and mutant PmGlmU and AGX1 uridylyltransferases.

PmGlmU and AGX1 had poor to no tolerance of substrates with bulkier alkynyl and diazirine functionalities (0-30% conversion, Figure 1C), which is in agreement with previous reports on the synthesis of UDP-GalNAc derivatives using AGX1.^31, 32^ UDP-sugars that cannot be accessed by enzymatic methods have instead been prepared through more lengthy chemical syntheses, most commonly by coupling an enzymatically prepared glycosyl-1-*O*-phosphate with UMP-morpholidate, a reaction that can take 3-5 days.^32, 37, 38^ Inspired by recent MOE reports where an F383A/G point mutation to AGX1 improved incorporation of GalNAz, GalNAlk, and GlcNDAz by increased biosynthetic production of the corresponding UDP-sugars,^15, 39–42^ we examined if analogous mutations to recombinant AGX1 and PmGlmU would improve OPME syntheses. Substrate promiscuity towards GalNAc derivatives in MOE labeling was enhanced with AGX1^F383A^ due to increased space in the catalytic pocket to accept bulkier C2-substituents.^41^ We therefore reasoned that mutating a similar position on PmGlmU, residue T199, may have a similar desired effect on this enzyme (Figure S9), and we recombinantly expressed the mutant enzymes PmGlmU^T199A^ and AGX1^F383A^ in *E. coli* (Supporting Information). While unfortunately no improvement in conversions was observed when PmGlmU^T199A^ was used in the OPME reactions with any substrate, AGX1^F383A^ had higher promiscuity with all substrates in comparison to the AGX1 wild type (WT) enzyme (Figure 1C). Notably, AGX1^F383A^ resulted in 40% and 55% conversion into UDP-GlcNAlk (**3**) and UDP-GlcNDAz (**4**), respectively, full conversion into UDP-GalNAlk (**7**), and 36% conversion into UDP-GalNDAz (**8**). Given that AGX1^F383A^ was observed to be the most promising enzyme for all the derivatives tested, it was used to synthesize all unnatural UDP-GlcNAc and UDP-GalNAc derivatives (**2-4** and **6-8**; Supporting Information) using an OPME approach, including compounds **7** and **8** which previously required chemical manipulations to access.^32^

Having prepared UDP-GlcNAc donor substrates **1-4**, we sought to assess the donor and acceptor substrate promiscuity of human glycosyltransferases B3GNT2 and B4GalT1, respectively, to access LacNAc derivatives. While several UDP-GlcNAc derivatives have been explored as donor substrates for bacterial β1,3-GlcNAcTs,^43^ many have not been examined as donors for B3GNT2. To date, UDP-GlcNDAz (**4**) has been assessed as a donor substrate for B3GNT2,^15^ and this enzyme has also been used to label asialofetuin with a C2-keto UDP-Glc donor derivative.^44^ Thus, we first examined each UDP-GlcNAc donor (**1-4**) as a substrate for B3GNT2 using a LacNAcβ-MU (**9**) disaccharide acceptor (Figure 2A). Conversion into trisaccharide products **10-13** was determined by HPLC-MS analysis (Figure 2B,D) after overnight incubation. As expected, the natural UDP-GlcNAc (**1**) substrate was the best donor (65% conversion). We were pleased to see that all modified GlcNAc substrates were tolerated and transfered by B3GNT2 (∼15-30% conversion). It is evident that as the size of the C2 *N*-acyl-functionality increases, transfer activity decreases: formation of the β1,3-GlcNAz-trisaccharide **11** proceeded with 26% conversion, whereas conversion into the β1,3-GlcNDAz-trisaccharide **13** was 17%. We were intrigued that the activity of B3GNT2 with UDP-GlcNDAz (**4**) was higher than a previous report by the Kohler group (17% *vs.* <1%).^15^ In their work, B3GNT2 activity was quantified using a UDP-Glo^TM^ glycosyltransferase assay after a 1 h reaction time. Thus, we hypothesize our increased conversion is likely attributed to an overnight (12-16 h) reaction time prior to quantification of trisaccharide conversion.

**Figure 2.**
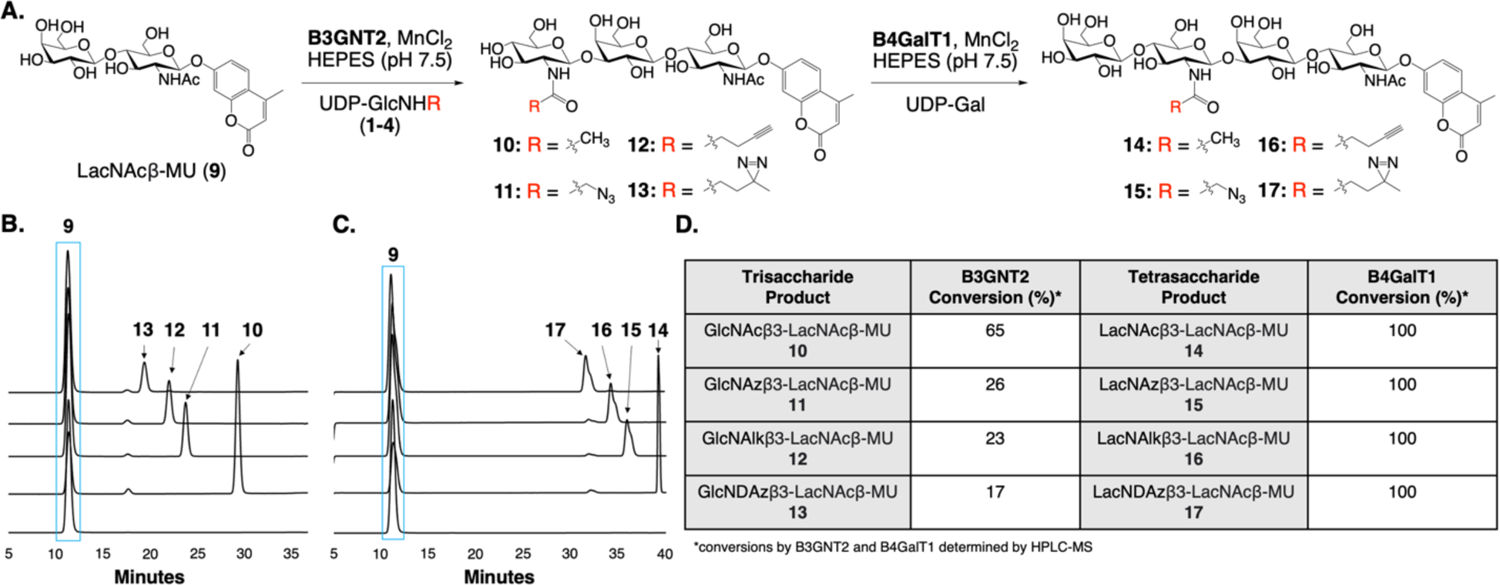
Substrate specificity of B3GNT2 and B4GalT1. (**A)** Enzymatic synthesis of diLacNAc glycoside derivatives using B3GNT2 and B4GalT1 starting from LacNAcβ-MU (**9**). (**B)** HPLC-MS analysis of B3GNT2 reactions to form trisaccharides **10-13** using acceptor **9** and UDP-GlcNAc derivatives **1-4**. (**C)** HPLC-MS analysis of B4GalT1 and UDP-Gal reactions to form diLacNAc tetrasaccharides **14-17** from trisaccharides **10-13**. (**D)** Reaction conversion percentages of B3GNT2 with UDP-GlcNAc derivatives **1-4,** and B4GalT1 and UDP-Gal with modified trisaccharide acceptors **10-13**.

Trisaccharides **10-13**, which contained different unnatural GlcNAc derivatives at the non-reducing terminus, were then assessed as acceptor substrates for β1,4-Gal addition with human B4GalT1 and UDP-Gal. Fortunately, after overnight incubation, all modified GlcNAc-glycosides (**10-13**) served as substrates for B4GalT1, and the formation of diLacNAc tetrasaccharide derivatives **14-17** proceeded with full conversion from the respective trisaccharide acceptor (Figure 2C,D). We were pleased to observe that human B4GalT1 tolerated all derivatives as bacterial and bovine β1,4-GalTs have been reported to have variable tolerance to modified GlcNAc-terminating acceptor substrates.^43, 45^ Our results are in alignment with crystal structures of the human B4GalT1 catalytic site showing that the C2 *N*-acyl group of the terminating GlcNAc on trisaccharide acceptors is relatively solvent-exposed, suggesting that this enzyme may be promiscuous towards substrates with modifications at this position.^15^

With the promising result that UDP-GlcNAz (**2**) is tolerated by B3GNT2, we next sought to glyco-engineer cells with GlcNAz and LacNAz motifs through exo-enzymatic glycan editing. We began by assessing the installation of GlcNAz onto a panel of cell lines, including Jurkat and U937 suspension cells, and HS-578-T and HEK293T adherent cells. Cells were treated for 2 hours concurrently with 0.2 mM UDP-GlcNAz (**2**), B3GNT2 and NanH, a sialidase from *Clostridium perfringens* which serves to desialylate cells and provide additional LacNAc acceptor sites to enhance B3GNT2 labeling (Figure 3A). Cell-surface GlcNAz introduction was detected by flow cytometry after a strain-promoted azide-alkyne cycloaddition (SPAAC) using a blue-fluorescent-DBCO conjugate, AZDye 405 DBCO (DBCO-405). Varying levels of GlcNAz incorporation was observed for each cell line after remodeling with B3GNT2 (Figure 3B), all of which were higher than the UDP-GlcNAz (**2**) or B3GNT2 alone controls (Figure S10). U937 and Jurkat cells had the lowest incorporation of GlcNAz, whereas HS-578-T and HEK293 cells showed a higher degree of labeling with HS-578-Ts labeling the strongest. We examined if GlcNAz incorporation onto Jurkat cells could be improved by increasing the concentration of UDP-GlcNAz (**2**), and found that labeling was concentration dependent and increasing the concentration of **2** from 0.2 mM to 2 mM enhanced labeling (Figure 3C).

**Figure 3.**
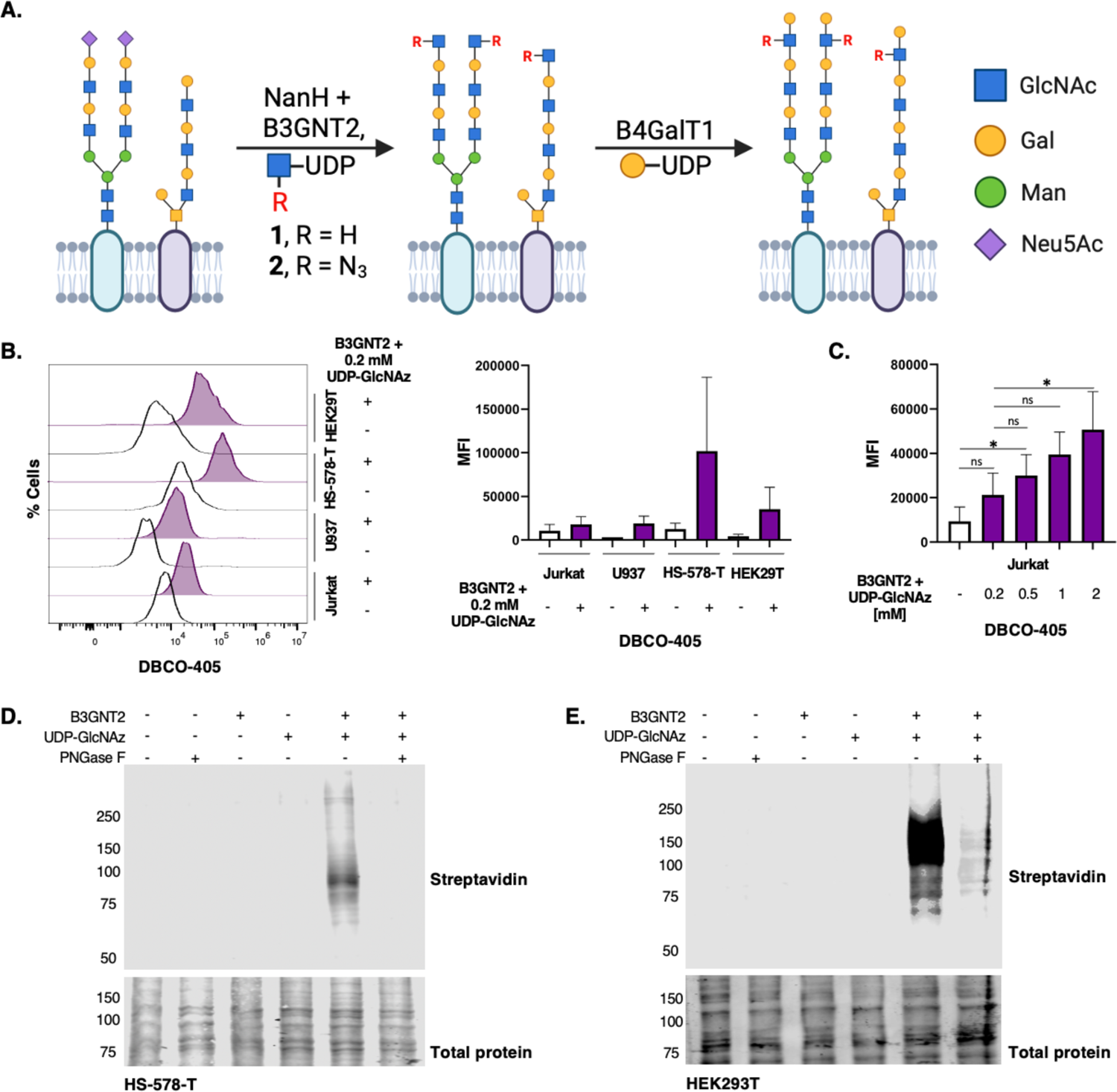
Cell-surface glycan remodeling with B3GNT2 and UDP-GlcNAz (**2**). **(A)** Cartoon representation of exo-enzymatic glycan editing strategy to label cells with GlcNAz and LacNAz motifs. **(B)** Flow cytometry analysis of GlcNAz-engineered cells. Cells were labeled with **2** (0.2 mM), B3GNT2 (250 mg/mL) and *C. perfringens* sialidase NanH (50 ug/mL) for 2 hours at 37°C. Cells were then conjugated to DBCO-405, co-stained with PI to exclude nonviable cells, and labeling with **2** on live-cell surfaces was assessed by flow cytometry. Median fluorescence intensity (MFI) of each sample was calculated. Error bars represent the standard deviation of two replicates (*n*=2). **(C)** Concentration-dependent labeling of live Jurkat cells with B3GNT2 and UDP-GlcNAz (**2**) assessed by flow cytometry after DBCO-405 conjugation. Error bars represent the standard deviation of three replicates (*n*=3). Statistical significance was assessed by unpaired *t*-test: \**p*<0.05; ns= not significant. **D,E)** Streptavidin Western blot analysis of GlcNAz-engineered cell lystastes. HS-578-T cells (**D)** or HEK293T cells (**E)** were conjugated to DBCO-biotin, lysed, incubated with or without PNGase F, and analyzed by immunoblot using streptavidin-800CW.

In addition to flow cytometry, cell-surface GlcNAz-labeled glycoproteins were analyzed by streptavidin Western blot after a SPAAC click reaction with DBCO-biotin. We initially examined HS-578-T cells, which showed the highest degree of labeling by flow cytometry (Figure 3B). Streptavidin immunoblot of GlcNAz-engineered HS-578-T lysates showed that a range of glycoproteins are labeled with B3GNT2 (Figure 3D). Interestingly, when GlcNAz-engineered HS-578-T lysates were treated with PNGase F, an enzyme that selectively cleaves *N*-glycans, a complete loss of streptavidin staining was observed, indicating that B3GNT2 was only labeling *N*-glycans on this cell line. This was unexpected as B3GNT2 is reported to act on LacNAc acceptors on *N*-glycans, *O*-glycans, and glycolipids.^25^ Since many cancer cell lines have truncated *O*-glycan structures,^46, 47^ we hypothesized that the apparent *N*-glycan specificity was attributed to the HS-578-T breast cancer cell line not displaying extended *O*-glycan structures that could serve as acceptor sites for B3GNT2. Thus, GlcNAz-engineered HEK293T cells were also analyzed by streptavidin Western blotting (Figure 3E). Similar to HS-578-T cells, a range of HEK293T cell glycoproteins were labeled with B3GNT2. After PNGase F treatment, a reduction in signal was observed with some streptavidin staining remaining on certain glycoproteins. These results indicate that GlcNAz installation on HEK293T cells occurred predominantly on *N*-glycans, and to a lesser extent on on *O*-glycans, a result in agreement with glycomics analysis of HEK293 cells reporting a low abundance of extended Core 2 *O*-glycans.^48^ Our data suggests that the apparent glycan sub-class specificity of glycosyltransferases employed in exo-enzymatic glycan editing is dependent on the cell line used, a factor that should be taken in to consideration when reporting enzyme specificity.

Having demonstrated that exo-enzymatic glyco-engineering could be used to install GlcNAz on multiple cell lines, we next sought to label cells with LacNAz epitopes using a two-step strategy. First, GlcNAz would be installed with B3GNT2 and UDP-GlcNAz (**2**), and subsequently a Gal residue would be transferred using B4GalT1 and UDP-Gal (Figure 3A). To assess the sequential build-up of LacNAz on cells, we used the lectin *Griffonia simplocifolia lectin-II* (GSL-II), which binds to terminal GlcNAc residues.^49^ A positive signal for GSL-II binding would be observed upon GlcNAz installation, and a loss of signal would be observed when the newly-installed GlcNAz is capped with Gal, forming a LacNAz motif. We assessed GSL-II binding to GlcNAz-engineered Jurkat cells labeled with a range of concentrations of UDP-GlcNAz (**2**) and compared this to exo-enzymatic glycan editing with natural UDP-GlcNAc (**1**) (Figure 4A-C). GlcNAz installation was detected by GSL-II binding above untreated control cells when 0.5 mM or greater UDP-GlcNAz (**2**) was used to engineer cells (Figure 4A, Figure S11), similar to results obtained with SPAAC (Figure 3C). Notably, when UDP-GlcNAc (**1**) was used to engineer Jurkat cells with B3GNT2, GSL-II binding was observed with 20-fold less donor substrate (0.025 mM UDP-GlcNAc). Thus, B3GNT2 is not as efficient at labeling cells with UDP-GlcNAz (**2**) in comparison to the natural UDP-GlcNAc donor substrate (Figure 4A), which is in alignment with our transfer assay using the LacNAcβ-MU (**9**) acceptor substrate (Figure 2). Indeed, exo-enzymatic labeling with 1 mM UDP-GlcNAz (**2**) had a similar GSL-II binding signal as when 0.025 mM UDP-GlcNAc (**1**) was used. While GSL-II binding could be substantially increased upon using higher concentrations of UDP-GlcNAc, a similar fold-increase was not observed when UDP-GlcNAz (**2**) was increased above 1 mM.

**Figure 4.**
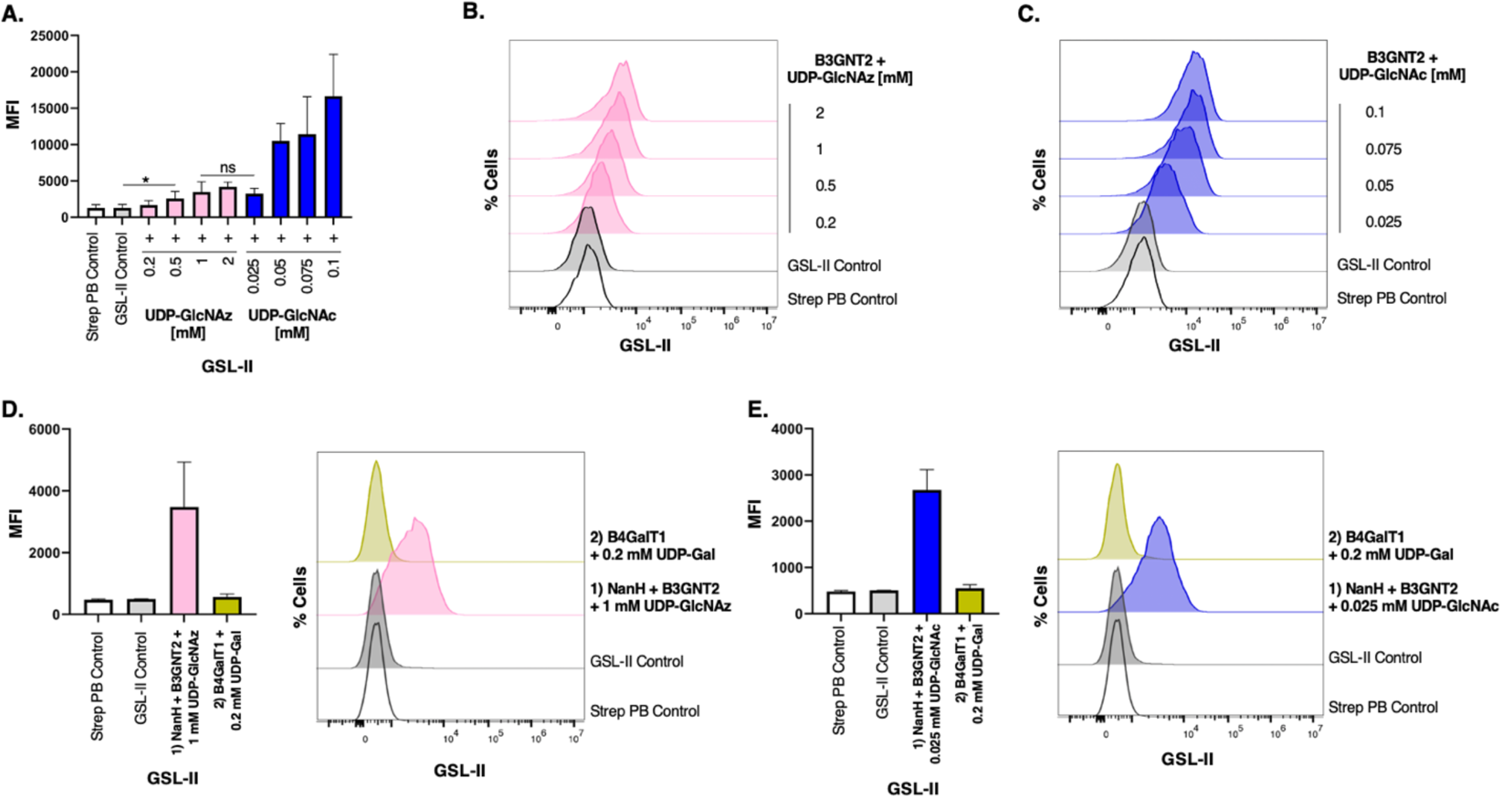
Cell-surface remodeling with LacNAz and LacNAc epitopes. **A-C)** GSL-II binding to GlcNAz- and GlcNAc-engineered Jurkat cells. Cells were treated with NanH, B3GNT2 and varying concentrations of UDP-GlcNAz (**2**) or UDP-GlcNAc (**1**) for 2 hours at 37°C. Cells were stained with biotinylated GSL-II, followed by Streptavidin-Pacific Blue and were co-stained with PI to exclude nonviable cells prior to flow cytometry analysis. **(A)** MFI of each sample was calculated. Error bars represent the standard deviation of three replicates (*n*=3). Statistical significance was assessed by unpaired *t*-test with \**p*<0.05, and ns= not statistically significant. Representative histograms of GSL-II binding are shown for **(B**) GlcNAz-engineered cells and **(C)** GlcNAc-engineered cells. **D,E)** LacNAz/LacNAc formation by B4GALT1 (250 μg/mL) and UDP-Gal (0.2 mM) treatment for 2 hours at 37°C on GlcNAz-engineered **(D)** and GlcNAc-engineered (**E**) cells. LacNAz/LacNAc formation was detected by flow cytometry analysis of GSL-II binding by observing a decrease in signal. MFI of each sample was calculated. Error bars represent the standard deviation of two replicates (*n*=2).

Given that 1 mM UDP-GlcNAz (**2**) and 0.025 mM UDP-GlcNAc (**1**) resulted in a similar degree of GlcNAz/GlcNAc installation based on GSL-II binding signals, these respective concentrations were used to display LacNAz and LacNAc motifs on cell-surfaces. Thus, after GlcNAz- or GlcNAc-engineering, cells were treated with B4GalT1 and UDP-Gal to cap the installed residues with Gal moieties. GSL-II binding data indicated that Gal residues were successfully transferred to the GlcNAz- and GlcNAc-engineered cells as a complete loss of binding was observed for both monosaccharides, indicating that all had been transformed into LacNAz or LacNAc, respectively (Figure 4D,E). These results are in agreement with our B4GalT1 acceptor substrate assay using UDP-Gal, which showed complete transfer of Gal to all modified GlcNAc acceptors (**10-13**) assessed (Figure 2).

In summary, we report an expanded scope for the OPME chemo-enzymatic synthesis of unnatural NT-sugars by employing mutant AGX1^F383A^. AGX1^F383A^ afforded access to GlcNAc- and GalNAc-based unnatural NT-sugars, and this enzyme could be used to synthesize derivatives previously not prepared through a fully enzymatic approach, including UDP-GalNAlk (**7**) and UDP-GalNDAz (**8**). The substrate promiscuity of recombinant human B3GNT2 and B4GalT1 using a LacNAcβ-MU glycoside (**9**) is also reported. While not as efficient as the native UDP-GlcNAc donor substrate, B3GNT2 can somewhat tolerate azide, alkyne and diazirine modifications at the C2 *N*-acyl position. In contrast, B4GalT1 could fully tolerate each of the derivatized GlcNAc acceptor substrates, demonstrating that diLacNAc functionalized with chemical reporters can be prepared by the sequential action of B3GNT2 and B4GalT1. Complimenting enzyme activity assays, we also performed proof-of-concept exo-enzymatic cell-surface glycan editing using B3GNT2 and B4GalT1. Successful GlcNAz installation on cells by B3GNT2 was confirmed by DBCO-405 *via* SPAAC and lectin staining with GSL-II. This cell-surface labeling strategy is amenable to multiple cell types, though differing levels of GlcNAz incorporation was observed between the cell lines employed. By immunoblotting two distinct cell line lysates treated with PNGase F, we confirm that B3GNT2 can label *N*- and *O*-glycans with suitable acceptor substrates. Our results highlight how multiple cell lines should be explored before confirming the specificity of glycosyltransferases used in exo-enzymatic glycan editing, particularly for enzymes that utilize LacNAc acceptors as many cancer cell lines do not biosynthesize extended *O*-glycan structures. Finally, we demonstrated the feasibility of glyco-engineering cell surfaces with LacNAz motifs by installing a Gal residue onto GlcNAz-engineered cells using B4GalT1 and UDP-Gal. While B3GNT2 has a lower tolerance for modified UDP-GlcNAc donor substrates, our work establishes that, in addition to monosaccharides like sialic acid and fucose, LacNAc glyco-motifs bearing bioorthogonal handles can be installed onto cell-surfaces using exo-enzymatic glycan editing. Strategies to improve the donor promiscuity of B3GNT2, such as bump-and-hole mutagenesis in the catalytic site, similar to mutant AGX1^F383A^,^42^ could afford mutant B3GNT2 enzymes that may enhance GlcNAc-derivative labeling, particularly with bulkier alkyne and diazirine analogues. Overall, this study lays the foundation for expanding the repertoire of glycosyltransferases and NT-sugar derivatives available for exo-enzymatic glycan-editing. It provides a strategy to install native and chemical-reporter functionalized LacNAcs onto cells, which have been challenging to selectively introduce, thus affording new opportunities to study the biological function of this glyco-motif.

## Associated Content

### Supporting Information

Supplementary figures of nucleotide-sugar synthesis, flow cytometry analysis, Western blot analysis; materials and methods for chemical synthesis, and biological procedures, and discussions of experimental and characterization details, ^1^H- and ^13^C-NMR spectra.

## Author Contributions

F.V.D.G and M.E.B. contributed equally as co-first authors and were intimately involved in the design of experiments, the generation of reagents, the execution of experiments, the interpretation of data, and the writing and editing of the manuscript. Order of authorship was determined based on the order of data presentation in the manuscript, both authors can list themselves first is their respective CV. F.V.D.G performed all chemical and enzymatic synthesis. M.E.B. performed all cell-surface exo-enzymatic glycan editing experiments. M.I.P. was involved in the chemical and enzymatic synthesis of NT-sugar analogues. C.J.C. oversaw the entirety of the project and was involved in developing the overall hypotheses being tested, the design of experiments, the interpretation of data, and the writing and editing of the manuscript and associated documents.

## Acknowledgements

The authors acknowledge the Natural Sciences and Engineering Research Council of Canada (NSERC), The New Frontiers in Research Fund (NFRF), The Canada Foundation for Innovation (CFI), GlycoNet and the Banting Research Foundation for funding supporting this work. M.E.B acknowledges NSERC for a CGS-D scholarship.

## References

1. Flynn, R. A.; Pedram, K.; Malaker, S. A.; Batista, P. J.; Smith, B. A. H.; Johnson, A. G.; George, B. M.; Majzoub, K.; Villalta, P. W.; Carette, J. E.;, et al. Small RNAs are modified with N-glycans and displayed on the surface of living cells. Cell, 2021, 184, 3109–3124.

2. Yang, W.; Fan, H.; Gao, X.; Gao, S.; Karnati, V. V. R.; Ni, W.; Hooks, W. B.; Carson, J.; Weston, B.; Wang, B. The First Fluorescent Diboronic Acid Sensor Specific for Hepatocellular Carcinoma Cells Expressing Sialyl Lewis X. Chem. Biol., 2004, 11, 439–448.

3. Pal, A.; Bérubé, M.; Hall, D. G. Design, Synthesis, and Screening of a Library of Peptidyl Bis(Boroxoles) as Oligosaccharide Receptors in Water: Identification of a Receptor for the Tumor Marker TF-Antigen Disaccharide. Angew. Chem., Int. Ed., 2010, 49, 1492–1495.

4. Thiemann, S.; Baum, L. G. Galectins and Immune Responses—Just How Do They Do Those Things They Do? Annu. Rev. Immunol., 2016, 34, 243–264.

5. Modenutti, C. P.; Capurro, J. I. B.; Di Lella, S.; Martí, M. A. The structural biology of galectin-ligand recognition: current advances in modeling tools, protein engineering, and inhibitor design. Front. Chem., 2019, 7, 823.

6. Girotti, M. R.; Salatino, M.; Dalotto-Moreno, T.; Rabinovich, G. A. Sweetening the hallmarks of cancer: Galectins as multifunctional mediators of tumor progression. J. Exp. Med., 2019, 217, e20182041.

7. Mkhikian, H.; Mortales, C. L.; Zhou, R. W.; Khachikyan, K.; Wu, G.; Haslam, S. M.; Kavarian, P.; Dell, A.; Demetriou, M. Golgi self-correction generates bioequivalent glycans to preserve cellular homeostasis. eLife, 2016, 5, e14814.

8. Yu, H.; Li, X.; Chen, M.; Zhang, F.; Liu, X.; Yu, J.; Zhong, Y.; Shu, J.; Chen, W.; Du, H. Integrated glycome strategy for characterization of aberrant LacNAc contained N-glycans associated with gastric carcinoma. Front. Oncol., 2019, 9, 636.

9. Ichikawa, T.; Nakayama, J.; Sakura, N.; Hashimoto, T.; Fukuda, M.; Fukuda, M. N.; Taki, T. Expression of N-acetyllactosamine and β1, 4-galactosyltransferase (β4GalT-I) during adenoma-carcinoma sequence in the human colorectum. J. Histochem. Cytochem., 1999, 47, 1593–1601.

10. Dube, D. H.; Bertozzi, C. R. Metabolic oligosaccharide engineering as a tool for glycobiology. Curr. Opin. Chem. Biol., 2003, 7, 616–625.

11. Laughlin, S. T.; Bertozzi, C. R. Imaging the glycome. Proc. Natl. Acad. Sci., 2009, 106, 12–17.

12. Zheng, X.; He, B.; Meng, L.; Xie, R. Recent Advances in Protein-Specific Glycan Imaging and Editing. ChemBioChem, 2023, 24, e202200778.

13. Kim, E. J. Chemical Reporters and Their Bioorthogonal Reactions for Labeling Protein O-GlcNAcylation. Molecules, 2018, 23, 2411.

14. Saha, A.; Bello, D.; Fernández-Tejada, A. Advances in chemical probing of protein O-GlcNAc glycosylation: structural role and molecular mechanisms. Chem. Soc. Rev., 2021, 50, 10451–10485.

15. Wu, H.; Shajahan, A.; Yang, J.-Y.; Capota, E.; Wands, A. M.; Arthur, C. M.; Stowell, S. R.; Moremen, K. W.; Azadi, P.; Kohler, J. J. A photo-cross-linking GlcNAc analog enables covalent capture of N-linked glycoprotein-binding partners on the cell surface. Cell. Chem. Biol., 2022, 29, 84–97.

16. Hart, G. W.; Housley, M. P.; Slawson, C. Cycling of O-linked β-N-acetylglucosamine on nucleocytoplasmic proteins. Nature, 2007, 446, 1017–1022.

17. Vocadlo, D. J.; Hang, H. C.; Kim, E.-J.; Hanover, J. A.; Bertozzi, C. R. A chemical approach for identifying O-GlcNAc-modified proteins in cells. Proc. Natl. Acad. Sci., 2003, 100, 9116–9121.

18. Hang, H. C.; Yu, C.; Kato, D. L.; Bertozzi, C. R. A metabolic labeling approach toward proteomic analysis of mucin-type O-linked glycosylation. Proc. Natl. Acad. Sci., 2003, 100, 14846–14851.

19. Boyce, M.; Carrico, I. S.; Ganguli, A. S.; Yu, S.-H.; Hangauer, M. J.; Hubbard, S. C.; Kohler, J. J.; Bertozzi, C. R. Metabolic cross-talk allows labeling of O-linked β-N-acetylglucosamine-modified proteins via the N-acetylgalactosamine salvage pathway. Proc. Natl. Acad. Sci., 2011, 108, 3141–3146.

20. Zhu, Y.; Wu, J.; Chen, X. Metabolic Labeling and Imaging of N-Linked Glycans in Arabidopsis Thaliana. *Angew. Chem.*, Int. Ed., 2016, 55, 9301–9305.

21. Batt, A. R.; Zaro, B. W.; Navarro, M. X.; Pratt, M. R. Metabolic Chemical Reporters of Glycans Exhibit Cell-Type-Selective Metabolism and Glycoprotein Labeling. ChemBioChem, 2017, 18, 1177–1182.

22. Nischan, N.; Kohler, J. J. Advances in cell surface glycoengineering reveal biological function. Glycobiology, 2016, 26, 789–796.

23. Griffin, M. E.; Hsieh-Wilson, L. C. Glycan engineering for cell and developmental biology. Cell. Chem. Biol., 2016, 23, 108–121.

24. Togayachi, A.; Sato, T.; Narimatsu, H. Comprehensive Enzymatic Characterization of Glycosyltransferases with a β3GT or β4GT Motif. In Methods in Enzymology, Vol. 416; Academic Press, 2006; pp 91–102.

25. Hao, Y.; Créquer-Grandhomme, A.; Javier, N.; Singh, A.; Chen, H.; Manzanillo, P.; Lo, M. C.; Huang, X. Structures and mechanism of human glycosyltransferase β1,3-N-acetylglucosaminyltransferase 2 (B3GNT2), an important player in immune homeostasis. J. Biol. Chem., 2021, 296, 100042.

26. Broszeit, F.; van Beek, R. J.; Unione, L.; Bestebroer, T. M.; Chapla, D.; Yang, J.-Y.; Moremen, K. W.; Herfst, S.; Fouchier, R. A.; de Vries, R. P. Glycan remodeled erythrocytes facilitate antigenic characterization of recent A/H3N2 influenza viruses. Nat. Commun., 2021, 12, 5449.

27. Wu, Z. L.; Person, A. D.; Anderson, M.; Burroughs, B.; Tatge, T.; Khatri, K.; Zou, Y.; Wang, L.; Geders, T.; Zaia, J.;, et al. Imaging specific cellular glycan structures using glycosyltransferases via click chemistry. Glycobiology, 2017, 28, 69–79.

28. Almaraz, R. T.; Li, Y. Labeling glycans on living cells by a chemoenzymatic glycoengineering approach. Biol. Open, 2017, 6, 923–927.

29. Yu, H.; Chen, X. One-pot multienzyme (OPME) systems for chemoenzymatic synthesis of carbohydrates. Org. Biomol. Chem., 2016, 14, 2809–2818.

30. Chen, Y.; Thon, V.; Li, Y.; Yu, H.; Ding, L.; Lau, K.; Qu, J.; Hie, L.; Chen, X. One-pot three-enzyme synthesis of UDP-GlcNAc derivatives. Chem. Commun., 2011, 47, 10815–10817.

31. Guan, W.; Cai, L.; Wang, P. G. Highly Efficient Synthesis of UDP-GalNAc/GlcNAc Analogues with Promiscuous Recombinant Human UDP-GalNAc Pyrophosphorylase AGX1. Chem. Eur. J., 2010, 16, 13343–13345.

32. Wen, L.; Gadi, M. R.; Zheng, Y.; Gibbons, C.; Kondengaden, S. M.; Zhang, J.; Wang, P. G. Chemoenzymatic Synthesis of Unnatural Nucleotide Sugars for Enzymatic Bioorthogonal Labeling. ACS Catal., 2018, 8, 7659–7666.

33. Bond, M. R.; Zhang, H.; Vu, P. D.; Kohler, J. J. Photocrosslinking of glycoconjugates using metabolically incorporated diazirine-containing sugars. Nat. Protoc., 2009, 4, 1044–1063.

34. Bannwarth, W.; Knorr, R. Formation of carboxamides with N,N,N′,N′-tetramethyl (succinimido) uronium tetrafluoroborate in aqueous / organic solvent systems. Tetrahedron Lett., 1991, 32, 1157–1160.

35. Gilormini, P. A.; Lion, C.; Vicogne, D.; Levade, T.; Potelle, S.; Mariller, C.; Guérardel, Y.; Biot, C.; Foulquier, F. A sequential bioorthogonal dual strategy: ManNAl and SiaNAl as distinct tools to unravel sialic acid metabolic pathways. Chem. Commun., 2016, 52, 2318–2321.

36. Li, Y.; Yu, H.; Chen, Y.; Lau, K.; Cai, L.; Cao, H.; Tiwari, V. K.; Qu, J.; Thon, V.; Wang, P. G.;, et al. Substrate Promiscuity of N-Acetylhexosamine 1-Kinases. Molecules, 2011, 16, 6396–6407.

37. Schultz, V. L.; Zhang, X.; Linkens, K.; Rimel, J.; Green, D. E.; Deangelis, P. L.; Linhardt, R. J. Chemoenzymatic Synthesis of 4-Fluoro-N-Acetylhexosamine Uridine Diphosphate Donors: Chain Terminators in Glycosaminoglycan Synthesis. J. Org. Chem., 2017, 82, 2243–2248.

38. Ahmadipour, S.; Miller, G. J. Recent advances in the chemical synthesis of sugar-nucleotides. Carbohydr. Res., 2017, 451, 95–109.

39. Yu, S.-H.; Boyce, M.; Wands, A. M.; Bond, M. R.; Bertozzi, C. R.; Kohler, J. J. Metabolic labeling enables selective photocrosslinking of O-GlcNAc-modified proteins to their binding partners. Proc. Natl. Acad. Sci., 2012, 109, 4834–4839.

40. Cioce, A.; Malaker, S. A.; Schumann, B. Generating orthogonal glycosyltransferase and nucleotide sugar pairs as next-generation glycobiology tools. Curr. Opin. Chem. Biol., 2021, 60, 66–78.

41. Cioce, A.; Bineva-Todd, G.; Agbay, A. J.; Choi, J.; Wood, T. M.; Debets, M. F.; Browne, W. M.; Douglas, H. L.; Roustan, C.; Tastan, O. Y.;, et al. Optimization of Metabolic Oligosaccharide Engineering with Ac4GalNAlk and Ac4GlcNAlk by an Engineered Pyrophosphorylase. ACS Chem. Biol., 2021, 16, 1961–1967.

42. Schumann, B.; Malaker, S. A.; Wisnovsky, S. P.; Debets, M. F.; Agbay, A. J.; Fernandez, D.; Wagner, L. J. S.; Lin, L.; Li, Z.; Choi, J.;, et al. Bump-and-Hole Engineering Identifies Specific Substrates of Glycosyltransferases in Living Cells. Mol. Cell, 2020, 78, 824–834.

43. Li, Y.; Xue, M.; Sheng, X.; Yu, H.; Zeng, J.; Thon, V.; Chen, Y.; Muthana, M. M.; Wang, P. G.; Chen, X. Donor substrate promiscuity of bacterial β1-3-N-acetylglucosaminyltransferases and acceptor substrate flexibility of β1-4-galactosyltransferases. Bioorg. Med. Chem., 2016, 24, 1696–1705.

44. Pasek, M.; Ramakrishnan, B.; Boeggeman, E.; Mercer, N.; Dulcey, A. E.; Griffiths, G. L.; Qasba, P. K. The N-acetyl-binding pocket of N-acetylglucosaminyltransferases also accommodates a sugar analog with a chemical handle at C2. Glycobiology, 2012, 22, 379–388.

45. Lau, K.; Thon, V.; Yu, H.; Ding, L.; Chen, Y.; Muthana, M. M.; Wong, D.; Huang, R.; Chen, X. Highly efficient chemoenzymatic synthesis of β1–4-linked galactosides with promiscuous bacterial β1–4-galactosyltransferases. Chem. Commun., 2010, 46, 6066–6068.

46. Rømer, T. B.; Aasted, M. K. M.; Dabelsteen, S.; Groen, A.; Schnabel, J.; Tan, E.; Pedersen, J. W.; Haue, A. D.; Wandall, H. H. Mapping of truncated O-glycans in cancers of epithelial and non-epithelial origin. Br. J. Cancer, 2021, 125, 1239–1250.

47. El-Schich, Z.; Zhang, Y.; Göransson, T.; Dizeyi, N.; Persson, J. L.; Johansson, E.; Caraballo, R.; Elofsson, M.; Shinde, S.; Sellergren, B. Sialic acid as a biomarker studied in breast cancer cell lines in vitro using fluorescent molecularly imprinted polymers. Appl. Sci., 2021, 11, 3256.

48. Huang, Y.-F.; Aoki, K.; Akase, S.; Ishihara, M.; Liu, Y.-S.; Yang, G.; Kizuka, Y.; Mizumoto, S.; Tiemeyer, M.; Gao, X.-D. Global mapping of glycosylation pathways in human-derived cells. Dev. Cell, 2021, 56, 1195–1209.

49. Bojar, D.; Meche, L.; Meng, G.; Eng, W.; Smith, D. F.; Cummings, R. D.; Mahal, L. K. A Useful Guide to Lectin Binding: Machine-Learning Directed Annotation of 57 Unique Lectin Specificities. ACS Chem. Biol., 2022, 2993–3012.

