## Supporting Information for "Glyco-Engineering Cell Surfaces by Exo-Enzymatic Installation of GlcNAz and LacNAz Motifs"

Fabiola V. De León González[§,1], Marie E. Boddington[§,2], Martha I. Prindl[1], Chantelle J. Capicciotti*[,1, 2, 3]
[1]Department of Chemistry, Queen's University, Kingston, K7L 3N6, Canada
[2]Department of Biomedical and Molecular Sciences, Queen's University, Kingston, K7L 3N6, Canada
[3]Department of Surgery, Queen's University, Kingston, K7L 3N6, Canada
§ These authors contributed equally to this work.

**Contents**

Supplemental Figures .......... **S3**
Chemical Methods .......... **S15**
Biological Methods .......... **S22**
NMR Characterization .......... **S26**
References .......... **S26**

#### Supplemental Figures

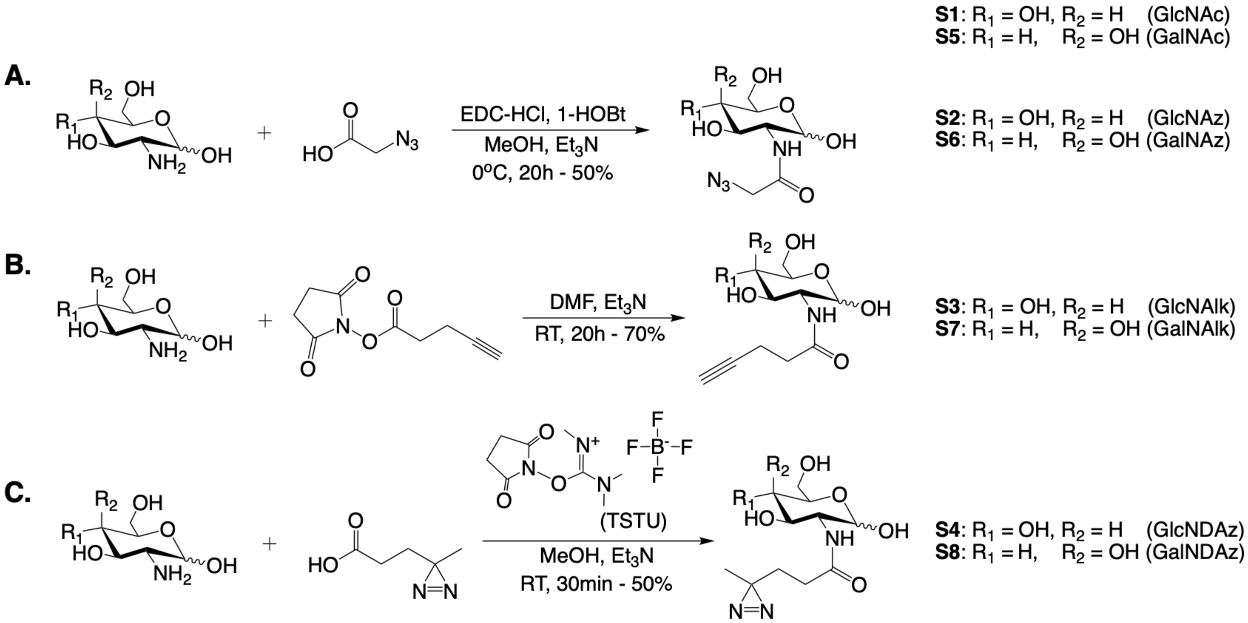


**Scheme S1. General synthesis of unnatural GlcNAc and GalNAc derivatives. (A)** Synthesis of azide-modified derivatives (**S2** & **S6**) by *in situ* EDC/HOBt-directed amide coupling. (**B)** Synthesis of alkyne-modified derivatives (**S3** & **S7**) by NHS-directed amide coupling.[2] **(C)** Synthesis of DAz-modified derivatives (**S4** & **S8**) by *in situ* TSTU-directed amide coupling.[3]

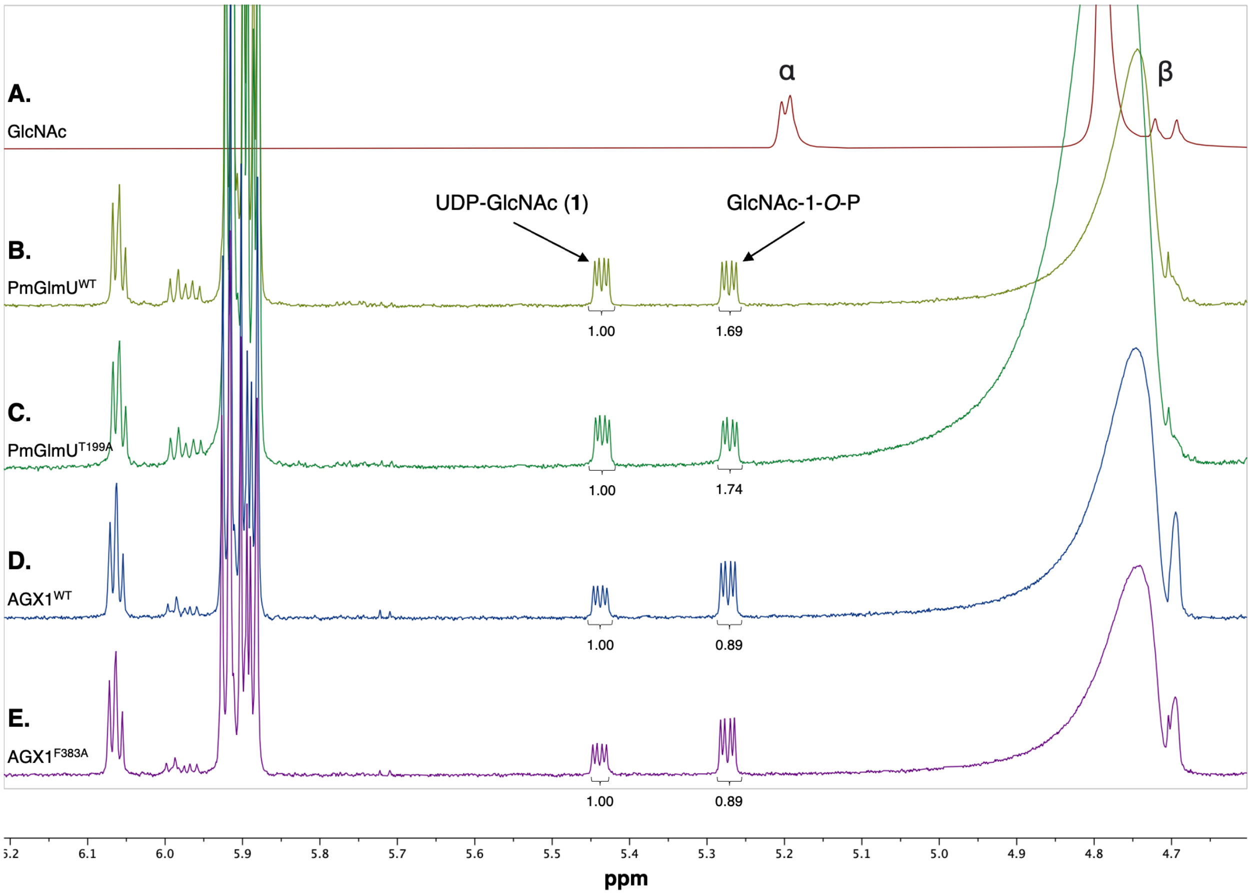


**Figure S1. Overlayed $^1$H-NMR (600 MHz; $D_2O$) spectra of crude OPME (NahK, Uridylyltransferase, & PmPpA) reaction synthesis of UDP-GlcNAc (1).** (**A**) Reference spectra of GlcNAc (**S1**) starting material. (**B**) OPME reaction using the bacterial PmGlmU$^{WT}$. **(C)** OPME reaction using the bacterial PmGlmU$^{T199A}$ mutant. (**D**) OPME reaction using the human AGX1$^{WT}$. **(E)** OPME reaction using the human AGX1$^{F383A}$ mutant. Anomeric proton shifts for **1** (5.44 ppm) and GlcNAc-1-*O*-P (5.27 ppm) used to calculate product conversions – integrations shown. Presaturation at 4.79 ppm.

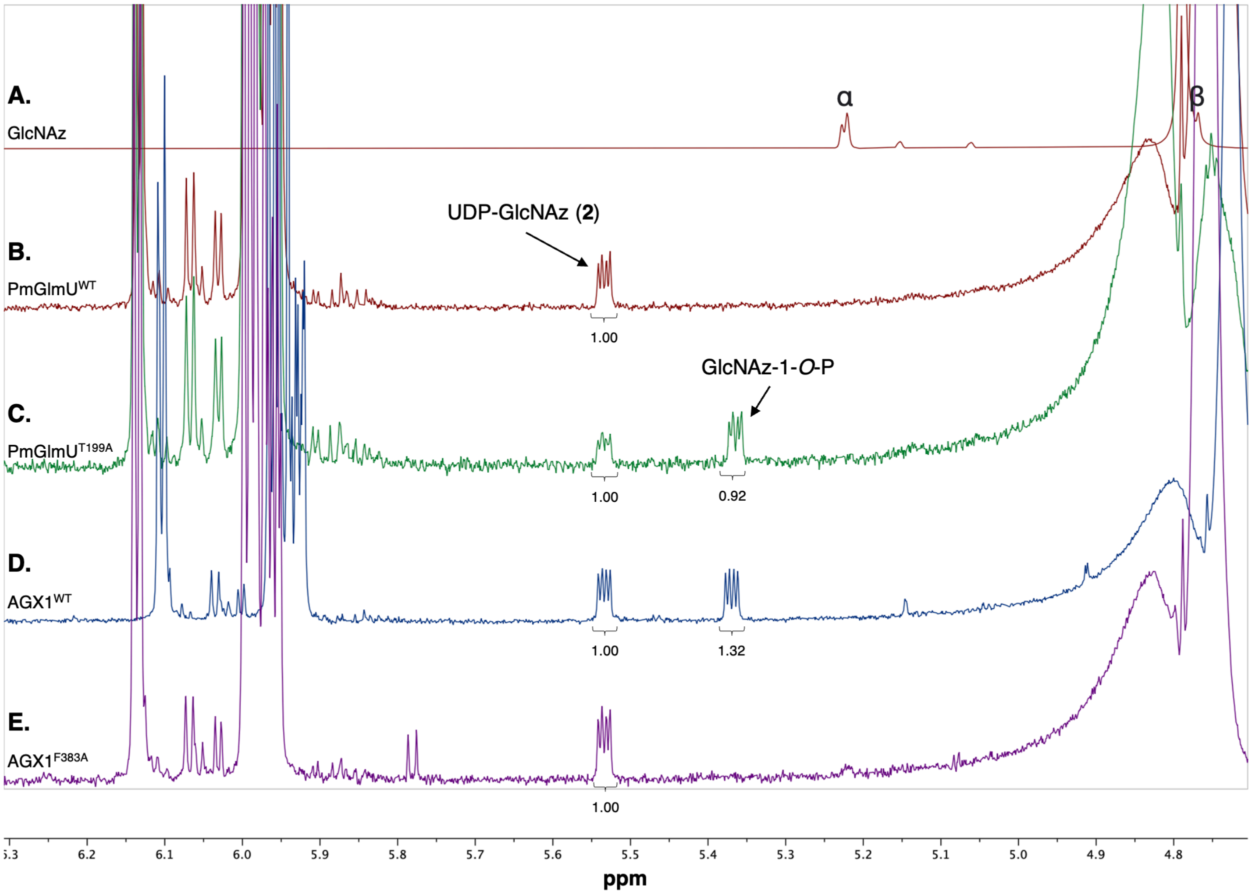


**Figure S2. Overlayed $^{1}$H-NMR (700 MHz; $D_2O$) spectra of crude OPME (NahK, Uridylyltransferase, & PmPpA) reaction synthesis of UDP-GlcNAz (2). (A).** Reference spectra of GlcNAz (**S2**) starting material. **(B)** OPME reaction using the bacterial PmGlmU$^{WT}$. **(C)** OPME reaction using the bacterial PmGlmU$^{T199A}$ mutant. **(D)** OPME reaction using the human AGX1$^{WT}$. **(E)** OPME reaction using the human AGX1$^{F383A}$ mutant. Anomeric proton shifts for **2** (5.53 ppm) and GlcNAz-1-*O*-P (5.37 ppm) used to calculate product conversions – integrations shown. Presaturation at 4.79 ppm.

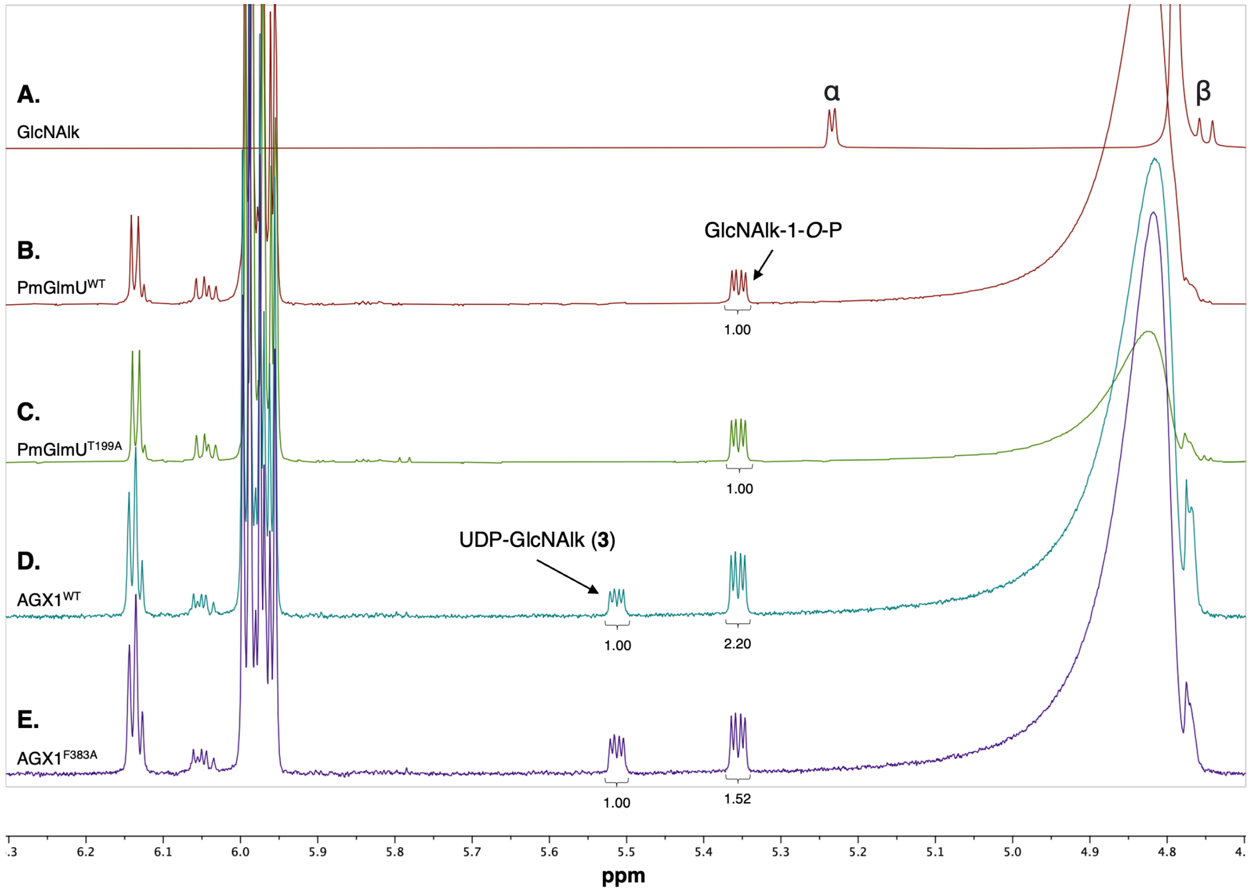


**Figure S3. Overlayed [1]H-NMR (600 MHz; $D_2O$) spectra of crude OPME (NahK, Uridylyltransferase, & PmPpA) reaction synthesis of UDP-GlcNAlk (3). (A).** Reference spectra of GlcNAlk (**S3**) starting material. **(B)** OPME reaction using the bacterial PmGlmU$^{WT}$. **(C)** OPME reaction using the bacterial PmGlmU$^{T199A}$ mutant. **(D)** OPME reaction using the human AGX1$^{WT}$. **(E)** OPME reaction using the human AGX1$^{F383A}$ mutant. Anomeric proton shifts for **3** (5.51 ppm) and GlcNAlk-1-*O*-P (5.35 ppm) used to calculate product conversions – integrations shown. Presaturation at 4.79 ppm.

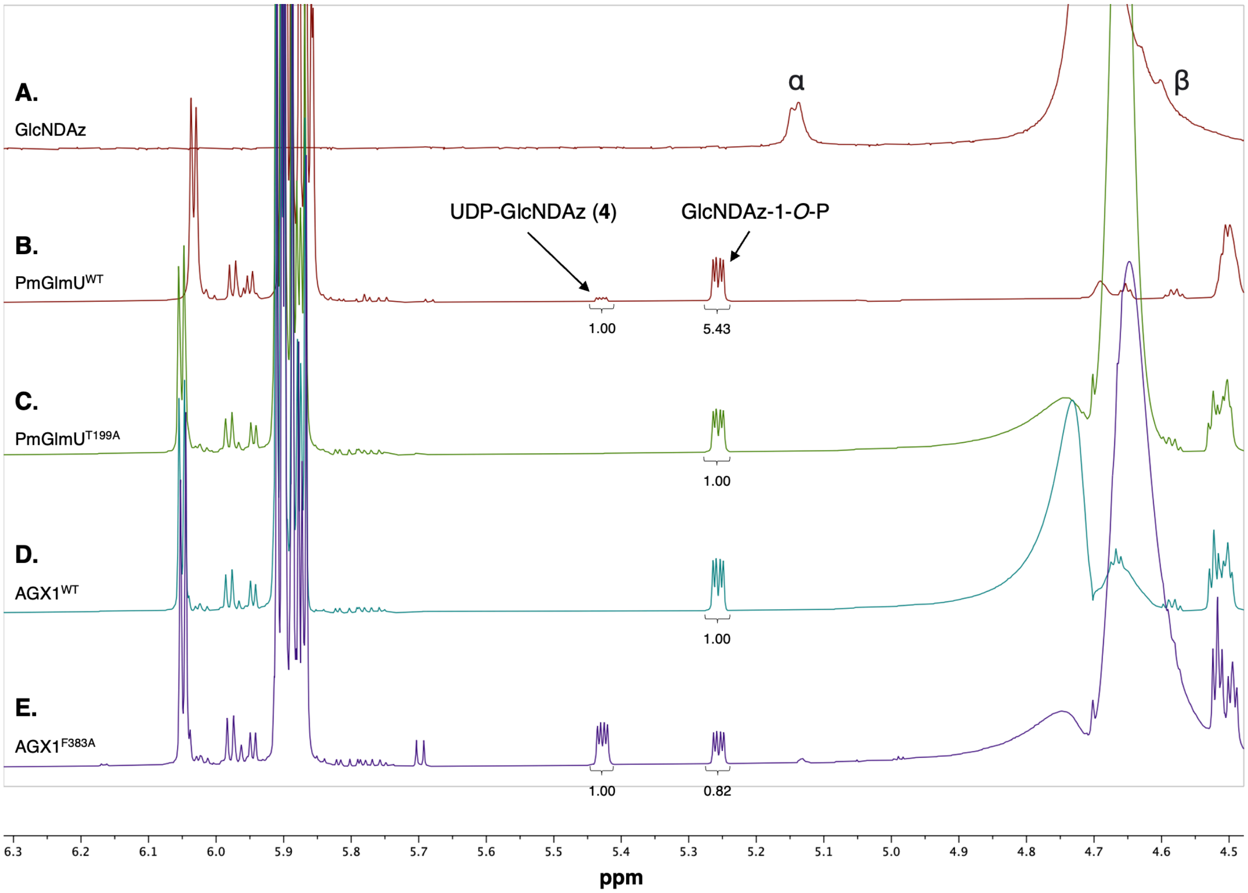


**Figure S4. Overlayed $^1$H-NMR (700 MHz; $D_2O$) spectra of crude OPME (NahK, Uridylyltransferase, & PmPpA) reaction synthesis of UDP-GlcNDAz (4). (A).** Reference spectra of GlcNDAz (**S4**) starting material. **(B)** OPME reaction using the bacterial PmGlmU$^{WT}$. **(C)** OPME reaction using the bacterial PmGlmU$^{T199A}$ mutant. **(D)** OPME reaction using the human AGX1$^{WT}$. **(E)** OPME reaction using the human AGX1$^{F383A}$ mutant. Anomeric proton shifts for **4** (5.43 ppm) and GlcNDAz-1-*O*-P (5.25 ppm) used to calculate product conversions – integrations shown. Presaturation at 4.79 ppm.

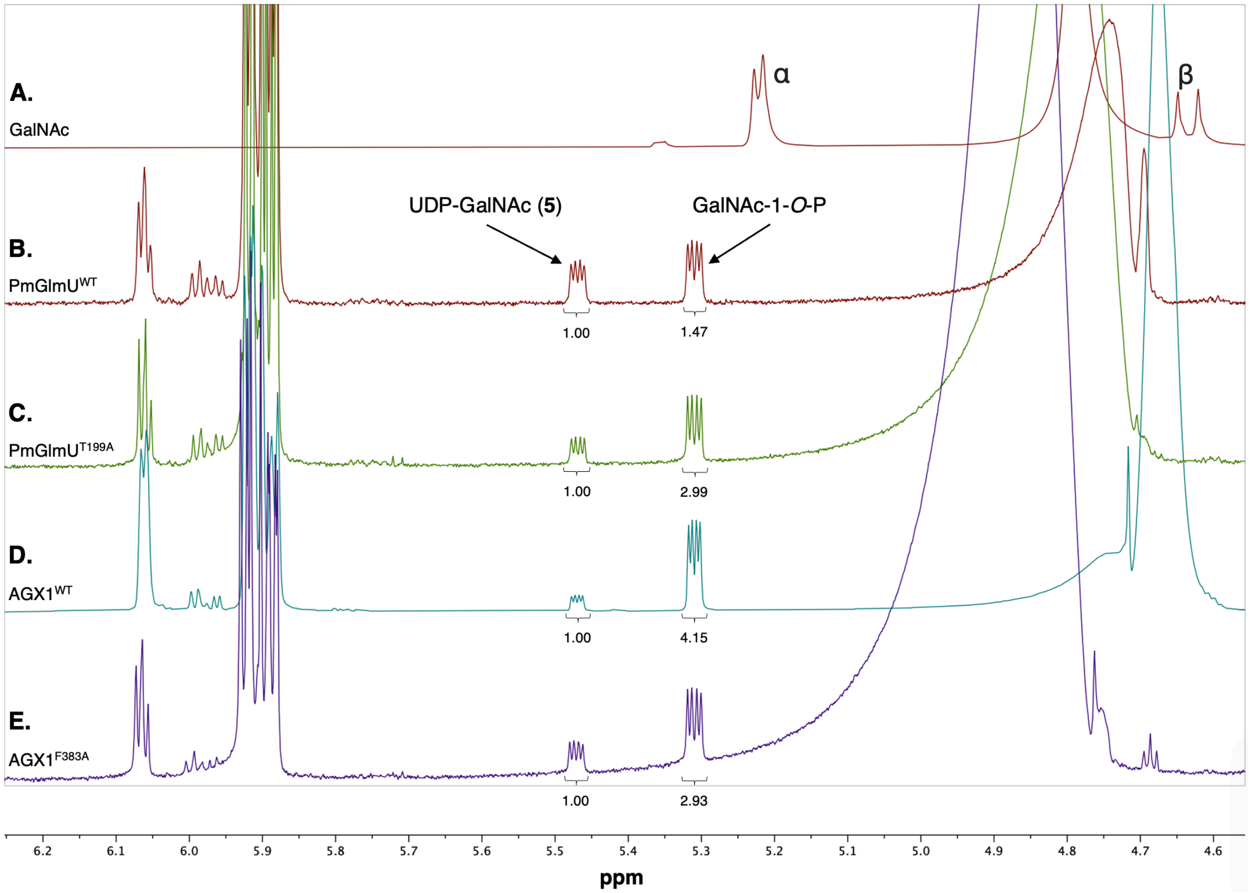


**Figure S5. Overlayed [1]H-NMR (600 MHz; $D_2O$) spectra of crude OPME (NahK, Uridylyltransferase, & PmPpA) reaction synthesis of UDP-GalNAc (5). (A).** Reference spectra of GalNAc (**S5**) starting material. **(B)** OPME reaction using the bacterial PmGlmU$^{WT}$. **(C)** OPME reaction using the bacterial PmGlmU$^{T199A}$ mutant. **(D)** OPME reaction using the human AGX1$^{WT}$. **(E)** OPME reaction using the human AGX1$^{F383A}$ mutant. Anomeric proton shifts for **5** (5.47 ppm) and GalNAc-1-*O*-P (5.31 ppm) used to calculate product conversions – integrations shown. Presaturation at 4.79 ppm.

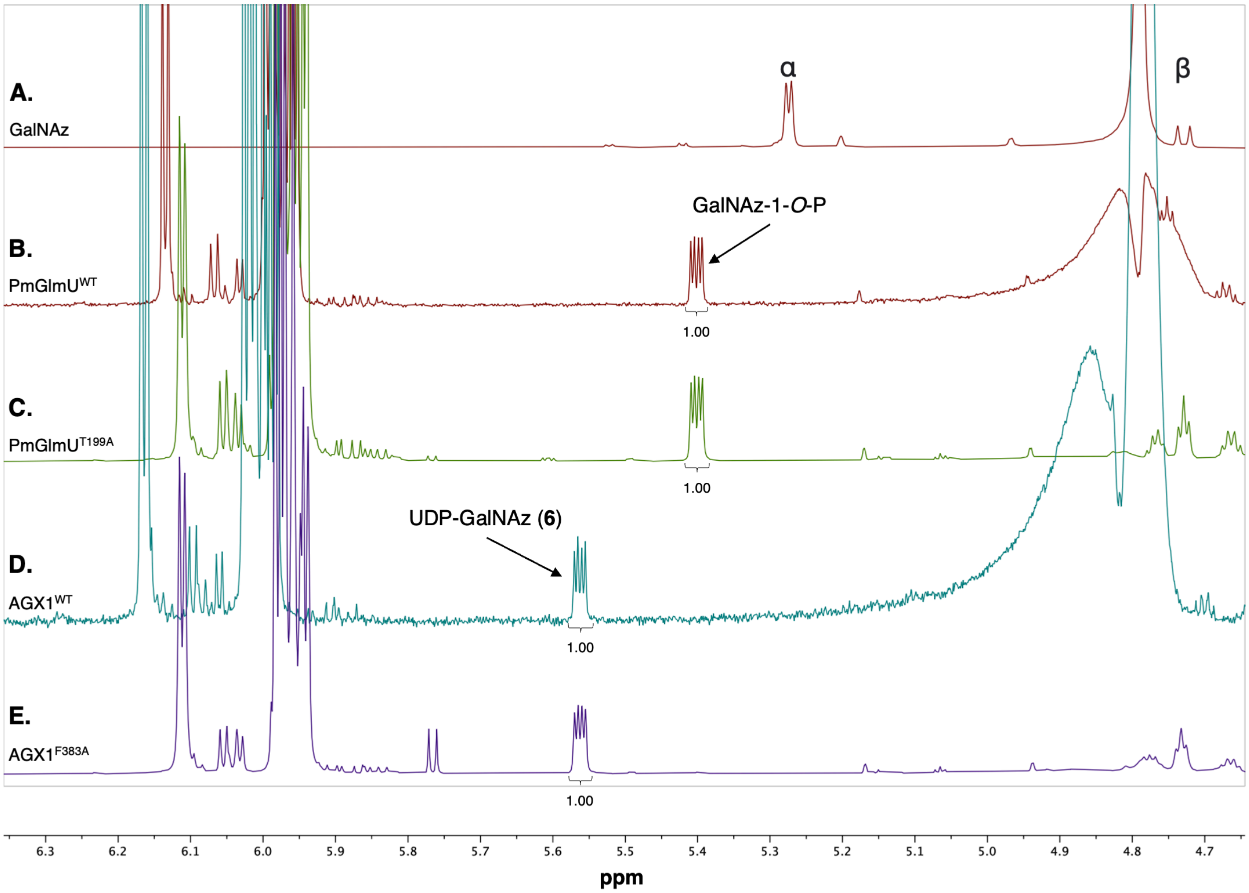


**Figure S6. Overlayed $^1$H-NMR (700 MHz; $D_2O$) spectra of crude OPME (NahK, Uridylyltransferase, & PmPpA) reaction synthesis of UDP-GalNAz (6). (A).** Reference spectra of GalNAz (**S6**) starting material. **(B)** OPME reaction using the bacterial PmGlmU$^{WT}$. **(C)** OPME reaction using the bacterial PmGlmU$^{T199A}$ mutant. **(D)** OPME reaction using the human AGX1$^{WT}$. **(E)**OPME reaction using the human AGX1$^{F383A}$ mutant. Anomeric proton shifts for **6** (5.56 ppm) and GalNAz-1-*O*-P (5.40 ppm) used to calculate product conversions – integrations shown. Presaturation at 4.79 ppm.

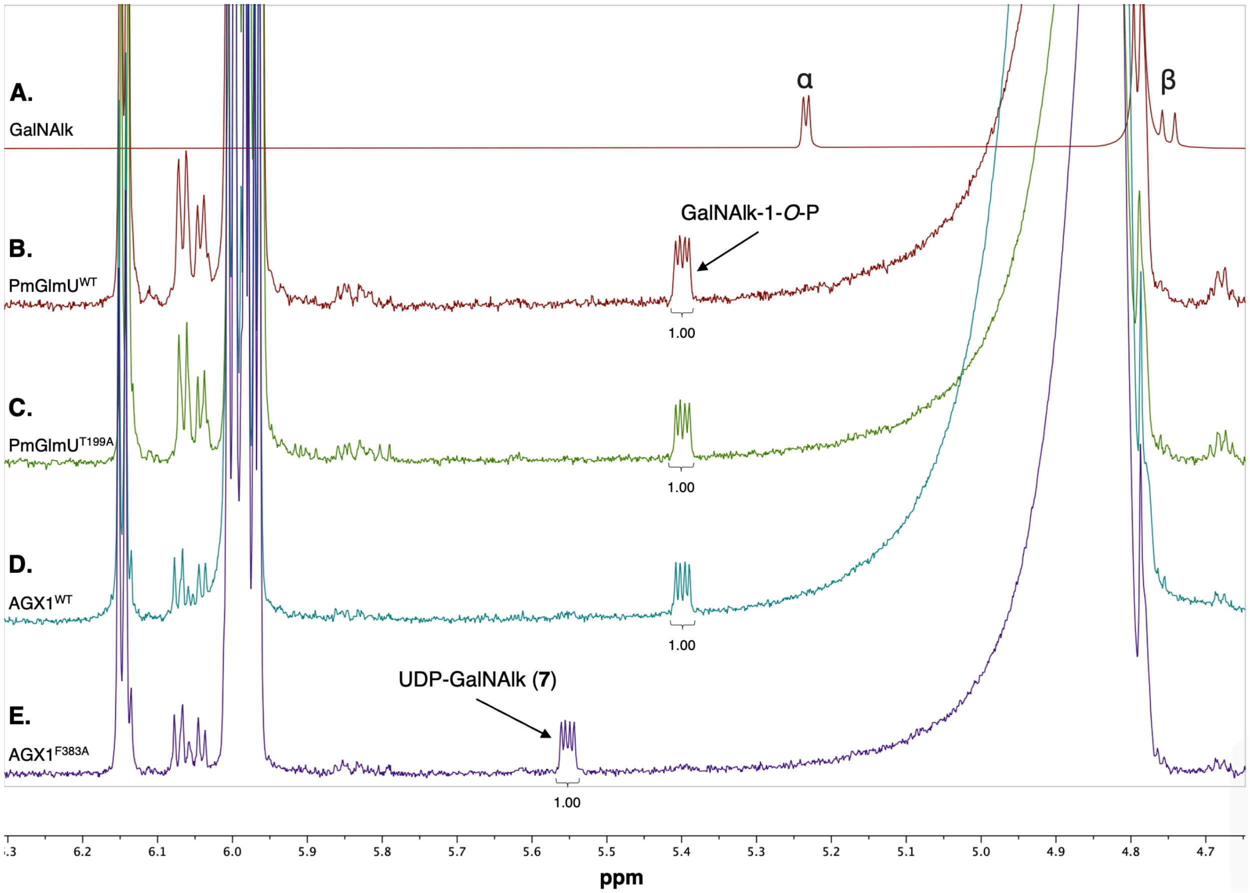


**Figure S7. Overlayed $^1$H-NMR (600 MHz; $D_2O$) spectra of crude OPME (NahK, Uridylyltransferase, & PmPpA) reaction synthesis of UDP-GalNAlk (7). (A).** Reference spectra of GalNAlk (**S7**) starting material. **(B)** OPME reaction using the bacterial PmGlmU$^{WT}$. **(C)** OPME reaction using the bacterial PmGlmU$^{T199A}$ mutant. **(D)** OPME reaction using the human AGX1$^{WT}$. **(E)** OPME reaction using the human AGX1$^{F383A}$ mutant. Anomeric proton shifts for **7** (5.55 ppm) and GalNAlk-1-*O*-P (5.40 ppm) used to calculate product conversions – integrations shown. Presaturation at 4.79 ppm.

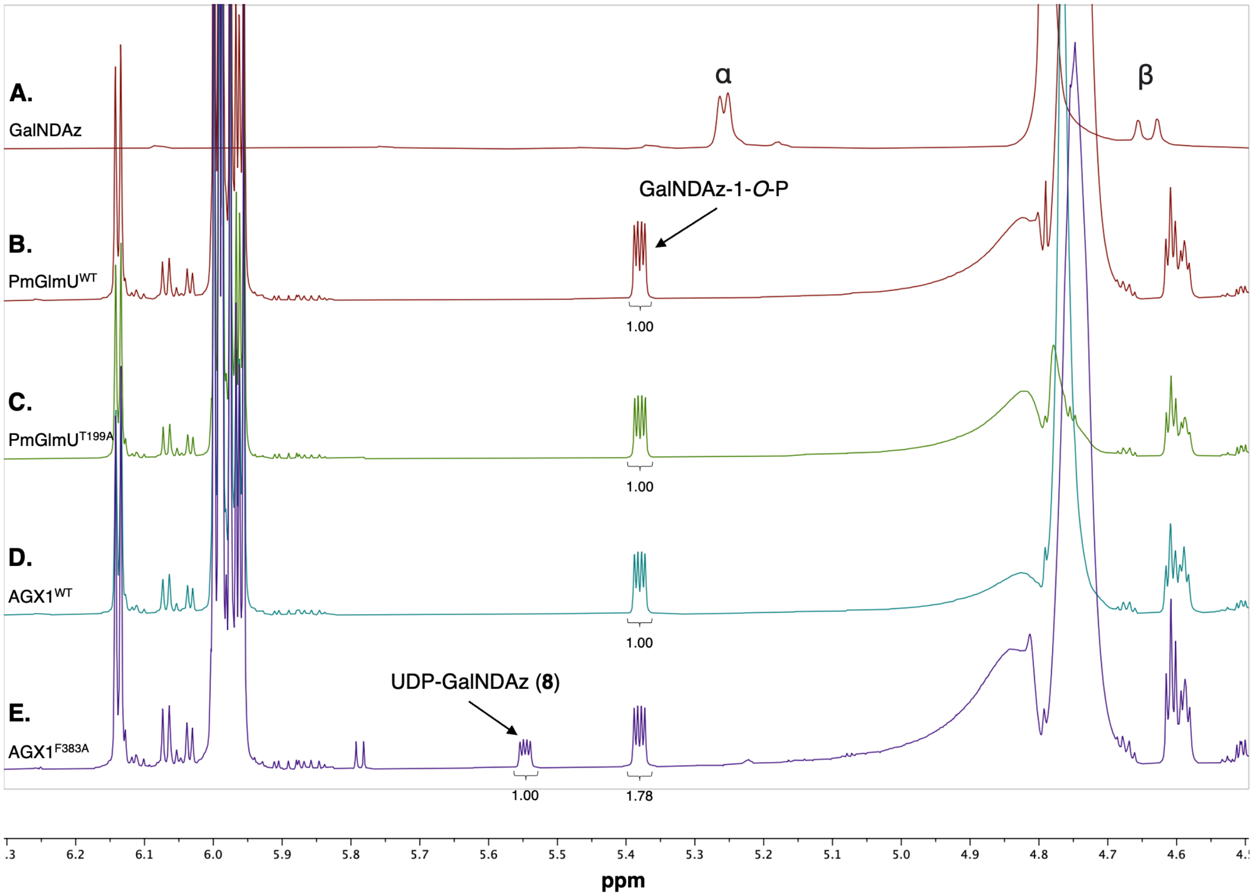


**Figure S8. Overlayed $^{1}$H-NMR (700 MHz; $D_2O$) spectra of crude OPME (NahK, Uridylyltransferase, & PmPpA) reaction synthesis of UDP-GalNDAz (8). (A).** Reference spectra of GalNDAz (**S8**) starting material. **(B)** OPME reaction using the bacterial PmGlmU$^{WT}$. **(C)** OPME reaction using the bacterial PmGlmU$^{T199A}$ mutant. **(D)** OPME reaction using the human AGX1$^{WT}$. **(E)** OPME reaction using the human AGX1$^{F383A}$ mutant. Anomeric proton shifts for **8** (5.55 ppm) and GalNDAz-1-*O*-P (5.38 ppm) used to calculate product conversions – integrations shown. Presaturation at 4.79 ppm.

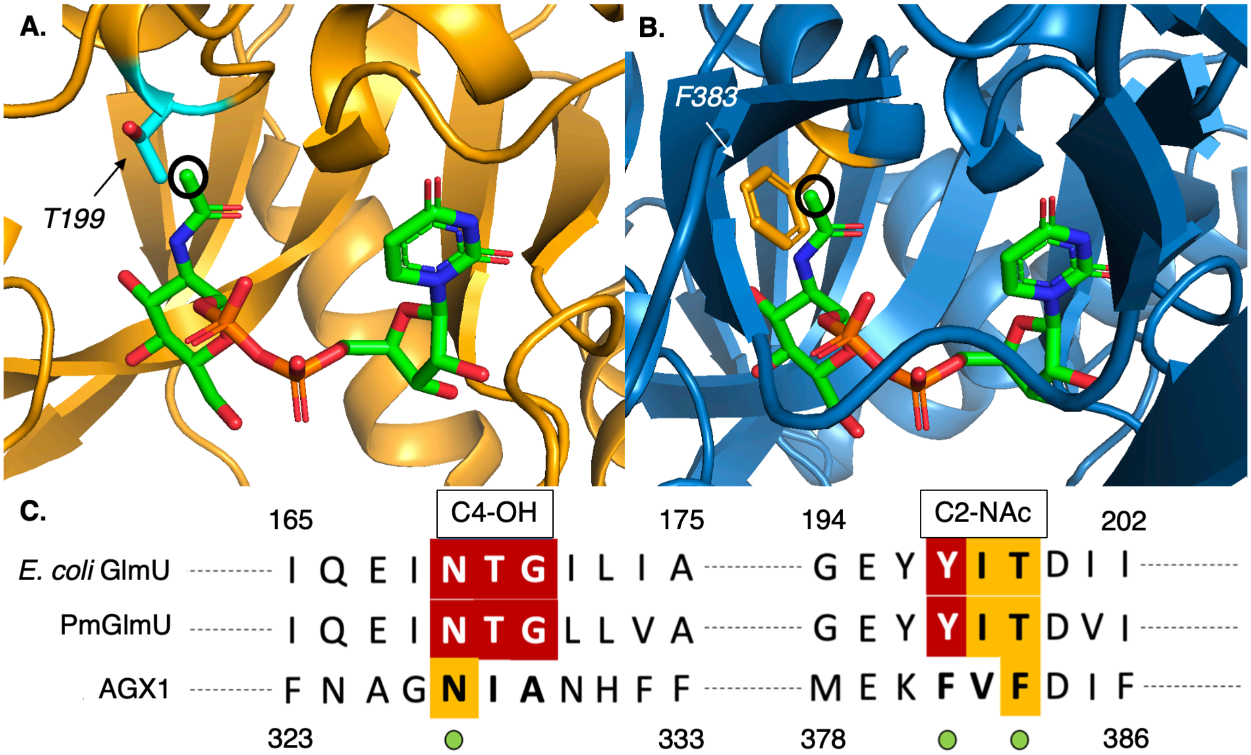


**Figure S9. Enzyme structure comparison study of PmGlmU and AGX1. (A)** The X-ray crystal structure of *E. coli* GlmU (PBD code 1FWY )[4] reveals that T199 is nearby the *N*-acetyl group of UDP-GlcNAc. **(B)** The X-ray crystal structure of human AGX1 (PDB code 1JV1)[5] reveals that F383 is nearby the *N*-acetyl group of UDP-GlcNAc; black circles indicate position where unnatural chemical modifications would be attached. **(C)** Partial amino acid sequence alignment of uridylyltransferases discussed in this work. Amino acid sequence comparison between EcGlmU and PmGlmU shows that the same residues are involved in nucleotide-sugar binding, with similar structural folds to AGX1 when bound to a UDP-GlcNAc substrate.[4, 5] Invariant amino acid residues in all sequences are highlight in white with a red background and conserved amino acid residues are shown in black with a yellow background; invariant residues for AGX1 are not reported in the literature. Amino acid residues involved in nucleotide-sugar recognition with the pyranose sugar are indicated by green circles and the specific pyranose position is indicated in the text above the residue.

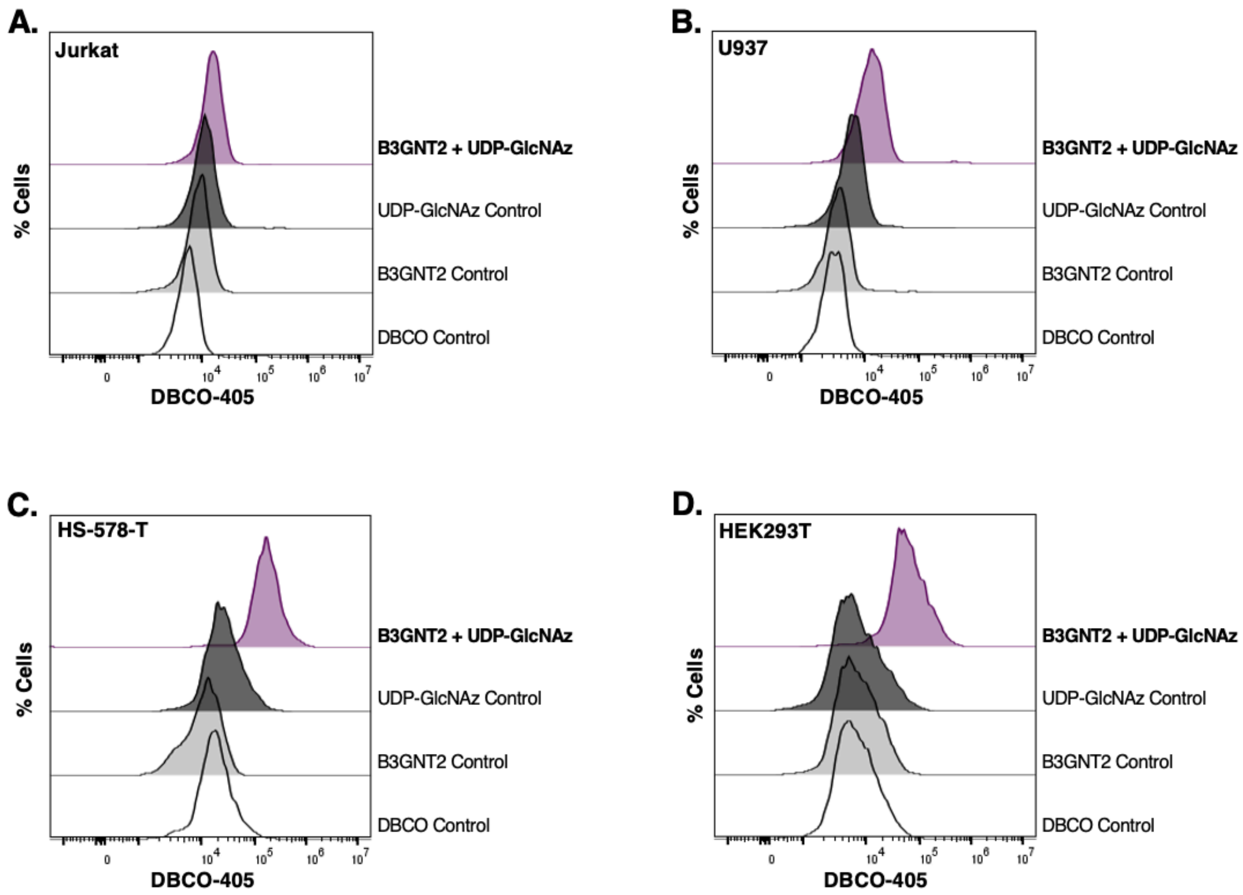


**Figure S10. Cell-surface glycan remodeling controls with B3GNT2 and UDP-GlcNAz** (**2**). Flow cytometry analysis of GlcNAz-engineered cells. Cells were stained with *C. perfringens* sialidase NanH (50 ug/mL), and varying conditions of UDP-GlcNAz **2** (0.2 or 0.5 mM) and B3GNT2 (250 mg/mL) for 2 hours at 37°C. Cells were then conjugated to DBCO-405, co-stained with PI to exclude nonviable cells, and labeling with **2** on live-cell surfaces was assessed by flow cytometry. The cell lines used were (**A**) Jurkat, **(B)** U937, **(C)** HS-578-T, and **(D)** HEK293T.

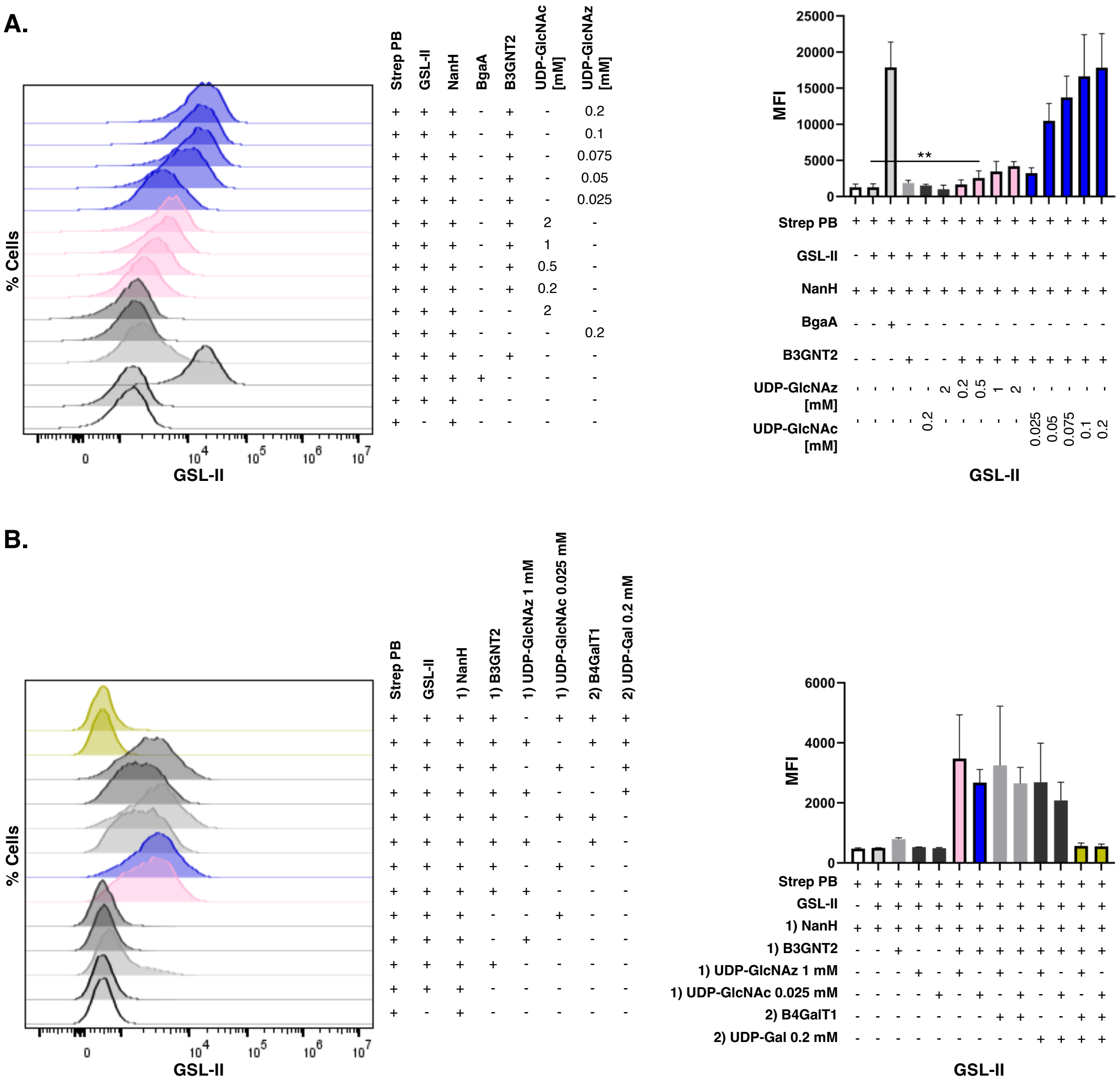


**Figure S11. Cell-surface glyco-engineering controls for LacNAz- and LacNAc-engineered cells. (A)** GSL-II binding to GlcNAc- and GlcNAz-engineered Jurkat cells. Cells were treated with NanH and varying conditions of B3GNT2, UDP-GlcNAc (**1**) or UDP-GlcNAz (**2**) for 2 hours at 37°C. Cells were stained with biotinylated GSL-II, followed by Streptavidin-Pacific Blue and were co-stained with PI to exclude nonviable cells prior to flow cytometry analysis. MFI of each sample was calculated. Error bars represent the standard deviation of three replicates ($n$=3). Statistical significance was assessed by unpaired $t$-test with $^{**}p<0.01$ **(B)** GSL-II binding to LacNAz/LacNAc engineered Jurkat cells. Cells were first treated with NanH and varying conditions of B3GNT2, UDP-GlcNAc (**1**) or UDP-GlcNAz (**2**) for 2 hours at 37°C. Next, GlcNAz- and GlcNAc-engineered cells were treated with or without B4GALT1 (250 µg/mL) and/or UDP-Gal (0.2 mM) for 2 hours at 37°C. Cells were stained with biotinylated GSL-II, followed by Streptavidin-Pacific Blue and were co-stained with PI to exclude nonviable cells prior to flow cytometry analysis. MFI of each sample was calculated. Error bars represent the standard deviation of two replicates ($n$=2).

#### Chemical Methods

##### Materials and general methods

All chemicals used for synthesis were purchased from Sigma Aldrich© unless otherwise noted: GlcNAcβ-MU, cat. M2133; UDP-Gal, cat. 670111-50MG. $GalNH_2$-HCl (cat. MG05030), GlcNAc (cat. MA00834), and GalNAc (cat. MA04390) were purchased from Biosynth©. TSTU (cat. CXZ041) and EDC-HCl (cat. CXZ005) were purchased from AAPPTec©. XBridge BEH amide columns (10x250mm & 4.6x250mm) and Sep-Pak® C18 1cc Vac cartridges (cat. 186000308) were purchased from Waters©. TLC analysis was performed with SiliaPlate Silica Gel 60-F254 (Silicycle) with detection by UV absorption (254 nm) and staining with para-anisaldehyde followed by heating. All moisture-sensitive reactions were carried out under an argon atmosphere. All chemical/enzymatic reactions involving the photo-crosslinkable diazirine (DAz) functional group performed in the dark to protect from degradation.

$^1$H and $^{13}$C NMR spectra were recorded on a Bruker© 500, 600, or 700 MHz spectrometer. Chemical shifts are reported in parts per million (ppm) relative to solvent residual peaks. NMR data are presented as follows: chemical shift, multiplicity (s = singlet, bs = broad singlet, d = doublet, dd = doublet of doublets, t = triplet, m = multiplet and/or multiple resonances), integration, coupling constant in Hz. All NMR signals were assigned using $^1$H-NMR, COSY, $^{13}$C-NMR, and HSQC experiments. High-resolution electrospray ionization (ESI) mass spectra were obtained using HPLC-MS (Agilent 1260 Infinity II LC-MSD System). Size exclusion chromatography was performed using a column packed with Bio-Gel® P-2 fine resins (Bio-Rad©, CA).

##### Chemical Synthesis

*N*-(4-Pentynoyloxy)succinimide (NHS-Alkyne):

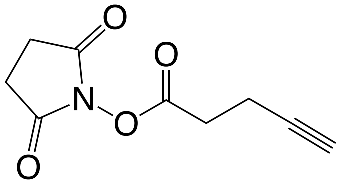


Synthesized by adapting previously reported protocol.[2] In a flame-dried reaction vessel under an argon atmosphere, 4-pentynoic acid (250 mg, 2 mmol), *N*-hydroxysuccinimide (NHS; 225mg, 2 mmol) and *N*-(3-dimethylaminopropyl)-*N'*-ethylcarbodiimide hydrochloride (EDC-HCl; 395 mg, 2 mmol) were dissolved in dry $CH_2Cl_2$ (4 mL). The reaction mixture was left to stir on ice for 20 hr while the temperature slowly increased to ambient level. The resulting mixture was diluted with $CH_2Cl_2$, extracted with $H_2O$ to remove the urea by-product, and the organic layer was dried over anhydrous $MgSO_4$ then concentrated *in vacuo*. The product was purified *via* silica column chromatography using a solvent system of 2:3 EtOAc/Hexanes to afford the NHS-Alkyne product as a white powder (100 mg, 20% isolated yield). Product was stored at -20 °C. Characterization data are consistent with that previously reported.[6] $^1$H NMR (500 MHz; $CDCl_3$): δ 2.86 (t, $J$ = 7.4 Hz, 2H), 2.82 (s, 4H), 2.59 (dddd, $J$ = 1.1, 2.7, 7.1, 8.3 Hz, 2H), 2.04 (t, $J$ = 2.6 Hz, 1H); $^{13}$C NMR (125 MHz; $CDCl_3$): δ 169.1, 167.1, 81.0, 70.1, 30.4, 25.7, 14.2. HRMS (ESI) *m/z* calculated for $C_9H_9NO_4$ (M+H) 196.0573, found 196.0596.

General amide coupling for the synthesis of azide-modified HexNAc derivatives (**Scheme S1A**):

Synthesized by adapting from previously reported protocol.[1] In a flame-dried round-bottom flask under an argon atmosphere, glucosamine-hydrochloride or galactosamine-hydrochloride was dissolved in minimal dry MeOH and triethylamine ($Et_3N$; 3 equiv.). Upon complete dissolution of reagents, 2-azidoacetic acid (1.5 equiv.)[7] was added. To this mixture, EDC-HCl (2 equiv.) and hydroxybenzotriazole (1-HOBt; 1 equiv.) were added and the reaction was left to stir on ice for 20 h while the temperature slowly increased to ambient level. The resulting reaction mixture was concentrated *in vacuo* and was acetylated by dissolving in pyridine (30 equiv.) and acetic anhydride (20 equiv.). The acetylation reaction was left to stir at room temperature overnight, and then was diluted in EtOAc; the organic layer was washed with 10% HCl, saturated sodium bicarbonate, and saturated NaCl. The organic layer was dried over anhydrous $MgSO_4$ then concentrated *in vacuo* and the crude mixture was purified *via* silica column chromatography using a solvent system of 2:3 EtOAc/Hexanes to afford the acetylated monosaccharide derivative as a white powder. The product was then dissolved in a minimal amount of MeOH, and NaOMe (1-3 drops) was added to deacetylate the sugar. The reaction was monitored by TLC (8:2 $CH_2Cl_2$/MeOH) and when the reaction was complete (~2-3 h), the pH of the mixture was adjusted to 7 using Amberlite® IR-120 H+ resin. The product was then purified on a P-2 Bio-Gel® column with 50 mM $NH_4HCO_3$ as the elution buffer to afford a pure mixture of anomers of the corresponding sugar.

General amide coupling synthesis of alkyne-modified HexNAc derivatives (**Scheme S1B)**:

Synthesized by adapting previously reported protocol.[2] In a flame-dried reaction vessel under an argon atmosphere, glucosamine-hydrochloride or galactosamine-hydrochloride was dissolved in minimal dry DMF and $Et_3N$ (3 equiv.). Upon complete dissolution of reagents, NHS-Alkyne (1.5 equiv.) was added and the resulting mixture was left to stir on ice for 20 h while the temperature slowly increased to ambient level. The reaction was monitored by TLC (8:2 $CH_2Cl_2$/MeOH) and upon completion the resulting reaction mixture was concentrated *in vacuo* and the crude was immediately purified *via* silica column chromatography using a gradient solvent system of 9:1-7:3 $CH_2Cl_2$/MeOH. Fractions containing product of interested were collected, concentrated *in vacuo* and further purified on a P-2 Bio-Gel® column with 50 mM $NH_4HCO_3$ as the elution buffer to afford a pure mixture of anomers of the corresponding product of interest.

General amide coupling synthesis of diazirine-modified HexNAc derivatives (**Scheme S1C**):

Synthesized by adapting previously reported protocol.[3] In a round-bottom flask, glucosamine-hydrochloride or galactosamine-hydrochloride, and *N, N, N', N'*-tetramethyl-*O*-(*N*-succinimidyl)-uronium tetrafluoroborate (TSTU; 1.5 equiv.) were dissolved in minimal 2:2:1 mixture of DMF/dioxane/$H_2O$ and *N,N*-diisopropylethylamine (DIPEA; 3 equiv.). Upon complete dissolution of reagents, 4,4-azo-pentanoic acid (diazirinie linker; 1.5 equiv.)[1] was added and the reaction was left to stir for 20-30 min. The reaction was monitored by TLC (8:2 $CH_2Cl_2$/MeOH) and upon completion the resulting reaction mixture was concentrated *in vacuo* and subsequently submerged in $CH_2Cl_2$. The tetramethylurea byproduct successfully dissolved in this solvent while the product of interested precipitated out of solution. The precipitate was collected by vacuum filtration and then further purified on a P-2 Bio-Gel® column with 50 mM $NH_4HCO_3$ as the elution buffer to afford a pure mixture of anomers of the corresponding sugar. All reaction products were stored at room temperature and protected from light-degradation.

2-Deoxy-2-azidoacetamido-D-glucopyranose (GlcNAz; **S2**):

From a 213 mg (0.99 mmol) scale reaction following the general protocol outlined for Scheme S1A, GlcNAz (**S2**) was obtained as a white powder (104 mg, 40% isolated yield). Product was stored at room temperature. Characterization data are consistent with that previously reported.[8] $^{1}$H NMR (500 MHz; $D_2O$) mixture of anomers: δ 5.23 (d, *J* = 3.5 Hz, 1H, Hα-1), 4.77 (d, *J* = 6.4 Hz, 1H, Hβ-1), 4.08 (s, 4H), 3.97-3.70 (m, 9H), 3.64-3.57 (m, 1H), 3.54-3.43 (m, 3H). $^{13}$C NMR (150 MHz; $D_2O$): δ 171.1, 170.7, 94.6, 90.8, 75.9, 73.6, 71.6, 70.6, 70.1, 69.9, 60.8, 60.6, 56.8, 54.2, 52.0, 51.8. HRMS (ESI) *m/z* calculated for $C_8H_{14}N_4O_6$ (M-H) 261.0827, found 261.0852.

2-Deoxy-2-(4-pentynoyl)amido-D-glucopyranose (GlcNAlk; **S3**):

From a 75 mg (0.29 mmol) scale reaction following the general protocol outlined for Scheme S1B, GlcNAlk (**S3**) was obtained as a white powder (65 mg, 72% isolated yield). Product was stored at room temperature. Characterization data are consistent with that previously reported.[9] $^{1}$H NMR (500 MHz; $D_2O$) mixture of anomers: δ 5.23 (d, *J* = 3.5 Hz, 1H, Hα-1), 4.75 (d, *J* = 8.5 Hz, 1H, Hβ-1), 3.98-3.84 (m, 4H), 3.83-3.69 (m, 4H), 3.60-3.46 (m, 3H), 3.26-3.19 (m, 2H), 2.91 (s, 1H). $^{13}$C NMR (125 MHz; $D_2O$): δ 175.2, 94.9, 90.9, 83.6, 75.9, 73.8, 71.6, 70.6, 70.2, 70.1, 69.6, 60.8, 60.6, 56.7, 54.1, 46.7, 42.7, 36.4, 34.7, 25.0, 14.4, 8.3. HRMS (ESI) *m/z* calculated for $C_{11}H_{17}NO_6$ (M-H) 258.1027, found 258.9951.

2-Deoxy-2-(4,4'-azo-pentanoyl)amido-D-glucopyranose (GlcNDAz; **S4**):

From a 84 mg (0.29 mmol) scale reaction following the general protocol outlined for Scheme S1C, GlcNDAz (**S4**) was obtained as a white powder (56 mg; 50% isolated yield). Product was stored at room temperature. Characterization data are consistent with that previously reported.[10] $^{1}$H NMR (500 MHz; $D_2O$) major anomer: δ 5.2 (d, *J* = 3.77 Hz, 1H, Hα-1), 3.94-3.61 (m, 6H), 3.58-3.38 (m, 2H), 2.30-2.13 (m, 2H), 1.78-1.59 (m, 2H), 1.0 (s, 3H). $^{13}$C NMR (125 MHz; $D_2O$): δ 175.9, 175.8, 94.9, 90.8, 75.9, 73.8, 71.5, 70.6, 70.1, 69.9, 60.7, 60.6, 56.7, 54.0, 30.4, 30.2, 29.9, 29.8, 26.2, 18.5. HRMS (ESI) *m/z* calculated for $C_{11}H_{19}N_3O_6$ (M-H) 288.1227, found 288.1212.

2-Deoxy-2-azidoacetamido-D-galactopyranose (GalNAz; **S6**):

From a 213 mg (0.99 mmol) scale reaction following the general protocol outlined for Scheme S1A, GalNAz (**S6**) was obtained as a white powder (88 mg, 34% isolated yield). Product was stored at room temperature. $^{1}$H NMR (500 MHz; $D_2O$) mixture of anomers: δ 5.25 (d, *J* = 3.75 Hz, 1H, Hα-1), 4.70 (d, *J* = 8.40 Hz, 1H, Hβ-1), 4.20 (dd, *J* = 3.7, 11.0,Hz, 1H), 4.15-4.09 (m, 2H), 4.07 (s, 4H), 4.01-3.90 (m, 4H), 3.81-3.68 (m, 6H). $^{13}$C NMR (150 MHz; $D_2O$): δ 171.2, 170.8, 95.1, 90.9, 75.2, 70.9, 70.6, 68.6, 67.9, 67.3, 61.2, 61.0, 53.9, 52.1, 51.8, 50.4. HRMS (ESI) *m/z* calculated for $C_8H_{14}N_4O_6$ (M-H) 261.0827, found 261.0851.

2-Deoxy-2-(4-pentynoyl)amido-D-galactopyranose (GalNAlk; **S7**):

From a 75 mg (0.29 mmol) scale reaction following the general protocol outlined for Scheme S1B, GalNAlk (**S7**) was obtained as a white powder (61 mg, 69% isolated yield). Product was stored at room temperature. $^{1}$H NMR (500 MHz; D2O) mixture of anomers: δ 5.23 (d, *J* = 3.6 Hz, 1H, Hα-1), 4.75 (d, *J* = 8.4 Hz, 1H, Hβ-1), 3.97-3.69 (m, 7H), 3.60-3.45 (m, 3H), 3.27-3.10 (m, 2H), 2.91 (s, 1H). $^{13}$C NMR (150 MHz; $D_2O$): δ175.3, 161.6, 95.4, 91.1, 75.2, 71.0, 70.6, 68.7, 67.9, 67.3, 62.6, 62.2, 61.3, 61.0, 57.0, 53.7, 50.3, 34.8, 34.4, 14.4. HRMS (ESI) *m/z* calculated for $C_{11}H_{17}NO_6$ (M-H) 258.1027, found 258.1000.

2-Deoxy-2-(4,4'-azo-pentanoyl)amido-D-galactopyranose (GalNDAz; **S8**):

From a 84 mg (0.29mmol) scale reaction following the general protocol outlined for Scheme S1C, GalNDAz (**S8**) was obtained as a white powder (50 mg, 45% isolated yield). Product was stored at room temperature. $^{1}$H NMR (500 MHz; $D_2O$) major anomer: δ 5.26 (d, *J* = 3.76 Hz, 1H, Hα-1), 4.17-4.01 (m, 2H), 3.94-3.61 (m, 6H), 4.00-3.63 (m, 6H), 2.27-2.15 (m, 2H), 1.74-1.58 (m, 2H), 1.0 (s, 3H). $^{13}$C NMR (150 MHz; $D_2O$): δ 176.2, 175.9, 112.2, 95.4, 91.0, 75.1, 71.1, 70.5, 68.7, 67.9, 67.3, 63.1, 61.2, 61.0, 56.3, 53.8, 50.3, 30.5, 30.3, 29.9, 29.8, 26.2, 18.5. HRMS (ESI) *m/z* calculated for $C_{11}H_{19}N_3O_6$ (M-H) 288.1227, found 288.1220.

**Chemo-Enzymatic Synthesis**

General enzymatic synthesis of UDP-GlcNAc (**1-4**) and UDP-GalNAc (**5-8**) derivatives:

The monosaccharide derivative (1.0 equiv.), ATP (1.2 equiv.), and UTP (5 equiv.) were dissolved in a Tris-HCl buffered solution (100 mM, pH 8.0) containing $MgCl_2$ (10 mM). To this solution, the enzymes *B. longum N*-acetylhexosamine-1-kinase (NahK; 15 mg/mmol substrate), the human uridylyltransferase mutant (AGX1$^{F383A}$; 20 mg/mmol substrate), and *P. multocida* strain Pm70 inorganic phosphatase (PmPpA; 10 mg/mmol substrate) were added. The reaction mixture was incubated overnight at 37 °C with gentle shaking. Reaction progress was monitored by TLC analysis (7:3 EtOH/1M $NH_4HCO_3$). Enzymes were removed by centrifugation using an Amicon® Ultra-30 centrifugal filter (10 kDa MWCO). The resulting filtrate was lyophilized and desalted on a P-2 Bio-Gel® column with 50 mM $NH_4HCO_3$ as the elution buffer to afford a mixture containing the product of interest, the glycosyl-1-*O*-phosphate synthetic intermediate (not observed for compounds **2**, **6**, or **7**), and a mixture of corresponding nucleosides and nucleotides. The mixture was redissolved in a Tris-HCl buffered solution (100 mM, pH 8.0) containing $CaCl_2$ (2 mM) and treated with an alkaline phosphatase (mAP; 20 mg/mmol substrate) to hydrolyze phosphate esters of nucleotides and the reaction mixture was subjected to size-exclusion purification by P-2 column to remove salts and glycerol prior to purification by HPLC-MS. After HPLC-MS purification, fractions containing desired product were combined, concentrated *in vacuo* and lyophilized to obtain the nucleotide-sugar product as a white solid.

Uridinyldiphosphate-α-2-deoxy-2-(amino)acetyl-D-glucopyranoside (UDP-GlcNAc; **1**)

For a 10 mg (45 μmol) scale reaction, UDP-GlcNAc (**1**) was obtained as a white powder (8.3 mg; 30% isolated yield). Product was stored at -80 °C. Characterization data are consistent with that previously reported.[11] $^{1}$H NMR (600 MHz, $D_2O$) δ 7.97 (d, *J* = 8.1 Hz, 1H), 6.02 – 5.96 (m, 2H), 5.53 (dd, *J* = 7.3, 3.3 Hz, 1H), 4.41 – 4.36 (m, 2H), 4.32-4.28 (m, 1H), 4.26 (ddd, *J* = 11.7, 4.6, 2.5 Hz, 1H), 4.20 (ddd, *J* = 11.8, 5.6, 3.0 Hz, 1H), 4.00 (dt, *J* = 10.5, 3.0 Hz, 1H), 3.94 (ddd, *J* = 10.1, 4.5, 2.4 Hz, 1H), 3.88 (dd, *J* = 12.5, 2.3 Hz, 1H), 3.84 – 3.80 (m, 2H), 3.59 – 3.54 (m, 1H), 2.09 (s, 3H). $^{13}$C NMR (151 MHz, $D_2O$) δ 174.79, 166.24, 151.84, 141.67, 102.68, 94.54, 88.50, 83.21, 73.80, 73.03, 70.98, 69.68, 69.54, 65.00, 60.36, 53.73, 22.10. HRMS (ESI) *m/z* calculated for $C_{17}H_{27}N_3O_{17}P_2$ (M-H) 606.0748, found 606.0743.

Uridinyldiphosphate-α-2-deoxy-2-azidoacetamido-D-glucopyranoside (UDP-GlcNAz; **2**):

For a 10 mg (38 μmol) scale reaction, UDP-GlcNAz (**2**) was obtained as a white powder (9.0 mg; 36% isolated yield). Product was stored at -80 °C. Characterization data are consistent with that previously reported.[11] $^{1}$H NMR (600 MHz; $D_2O$): δ 7.96 (d, *J* = 8.1 Hz, 1H), 5.98 (m, 2H), 5.55 (dd, *J* = 7.2, 3.4 Hz, 1H), 4.41-4.35 (m, 2H), 4.31-4.23 (m, 2H), 4.22-4.17 (m, 1H), 4.14 (s, 1H), 4.10-4.05 (m, 2H), 3.97-3.93 (m, 1H), 3.90-3.80 (m, 3H), 3.71-3.63 (m, 1H), 3.58 (t, *J* = 9.6

Hz, 1H). $^{13}C$ NMR (151 MHz, $D_2O$) δ 170.96, 166.49, 152.01, 141.67, 102.69, 94.36, 88.65, 83.18, 73.76, 73.08, 70.88, 69.66, 69.51, 65.04, 60.36, 51.63. HRMS (ESI) *m/z* calculated for $C_{17}H_{26}N_6O_{17}P_2$ (M-H) 647.0762, found 647.0757.

Uridinyldiphosphate-α-2-deoxy-2-(4-pentynoyl)amido-D-glucopyranoside (UDP-GlcNAlk; **3**):

For a 10 mg (39 μmol) scale reaction, UDP-GlcNAlk (**3**) was obtained as a white powder (7.0 mg; 28% isolated yield). Product was stored at -80 °C. Characterization data are consistent with that previously reported.[12] $^1H$ NMR (600 MHz; $D_2O$): δ 7.98 (d, $J$ = 8.1 Hz, 1H), 5.99 (m, 2H), 5.53 (dd, $J$ = 7.2, 3.4 Hz, 1H), 4.41-4.36 (m, 2H), 4.32-4.17 (m, 3H), 4.04 (dt, $J$ = 10.6, 3.1 Hz, 1H), 3.97-3.93 (m, 1H), 3.91-3.87 (m, 2H), 3.85-3.80 (m, 2H), 3.57 (t, $J$ = 9.5 Hz, 1H), 2.49-2.64 (m, 5H), 2.38 (t, $J$ = 2.55 Hz, 1H). $^{13}C$ NMR (150 MHz; $D_2O$): δ175.19, 166.27, 151.84, 141.69, 102.70, 94.67, 88.52, 83.71, 83.26, 83.20, 73.82, 73.04, 70.93, 70.02, 69.67, 69.57, 65.01, 60.35, 53.66, 53.61, 34.31, 14.34. HRMS (ESI) *m/z* calculated for $C_{20}H_{29}N_3O_{17}P_2$ (M-H) 644.0904, found 644.0911.

Uridinyldiphosphate-α-2-deoxy-2-(4,4'-azo-pentanoyl)amido-D-glucopyranoside (UDP-GlcNDAz; **4**):

For a 10 mg (35 μmol) scale reaction, UDP-GlcNDAz (**4**) was obtained as a white powder (6.3 mg; 27% isolated yield). Product was stored at -80 °C. Characterization data are consistent with that previously reported.[12] $^1H$ NMR (600 MHz; $D_2O$): δ 7.99 (d, $J$ = 8.1 Hz, 1H), 6.00 (m, 2H), 5.53 (dd, $J$ = 7.1, 3.3 Hz, 1H), 4.42-4.36 (m, 2H), 4.32-4.18 (m, 3H), 4.02-4.00 (m, 2H ), 3.97-3.93 (m, 1H), 3.91-3.86 (m, 1H), 3.85-3.80 (m, 2H), 3.57 (t, $J$ = 9.6 Hz, 1H), 2.37-2.25 (m, 2H), 1.76-1.63 (m, 2H), 1.04 (s, 3H). $^{13}C$ NMR (150 MHz; $D_2O$): δ 175.94, 166.24, 151.83, 141.73, 102.68, 94.61, 88.55, 83.25, 83.19, 73.81, 73.02, 70.95, 69.66, 69.59, 65.00, 60.34, 59.25, 53.60, 30.13, 29.84, 26.31, 18.53. HRMS (ESI) *m/z* calculated for $C_{20}H_{31}N_5O_{17}P_2$ (M-H) 674.1122, found 674.1125.

Uridinyldiphosphate-α-2-deoxy-2-(amino)acetyl-D-galactopyranoside (UDP-GalNAc; **5**):

For a 10 mg (45 μmol) scale reaction, UDP-GalNAc (**5**) was obtained as a white powder (6.9 mg; 25% isolated yield). Product was stored at -80 °C. Characterization data are consistent with that previously reported.[13] $^1H$ NMR (600 MHz; $D_2O$): δ 7.97 (d, $J$ = 8.2 Hz, 1H), 5.99 (m, 2H), 5.56 (dd, $J$ = 7.3, 3.4 Hz, 1H), 4.41-4.36 (m, 2H), 4.32-4.18 (m, 5H), 4.06 (d, $J$ = 2.7 Hz, 1H), 3.98 (dd, $J$ = 11, 3.2 Hz, 1H), 3.83-3.72 (m, 2H), 2.10 (s, 1H). $^{13}C$ NMR (150 MHz; $D_2O$): δ 174.97, 166.22, 151.80, 141.65, 102.63, 94.71, 88.44, 83.25, 83.20, 73.77, 72.04, 69.65, 68.37, 67.65, 64.98, 61.03, 49.77, 22.13. HRMS (ESI) *m/z* calculated for $C_{17}H_{27}N_3O_{17}P_2$ (M-H) 606.0748, found 606.0746.

Uridinyldiphosphate-α-2-deoxy-2-azidoacetamido-D-galactopyranoside (UDP-GalNAz; **6**):

For a 10 mg (38 μmol) scale reaction, UDP-GalNAz (**6**) was obtained as a white powder (5.7 mg; 23% isolated yield). Product was stored at -80 °C. Characterization data are consistent with that previously reported.[13] $^1H$ NMR (600 MHz; $D_2O$): δ 7.97 (d, $J$ = 8.1 Hz, 1H), 5.99 (m, 2H), 5.58 (dd, $J$ = 7.2, 3.5 Hz, 1H), 4.42-4.35 (m, 2H), 4.35-4.28 (m, 2H), 4.28-4.23 (m, 1H), 4.23-4.18 (m, 2H), 4.15 (s, 1H), 4.09 (s, 1H), 4.07-4.05 (m, 1H), 4.02-3.96 (dd, $J$ = 10.9, 3.2 Hz, 1H), 3.83-3.73 (m, 2H). $^{13}C$ NMR (150 MHz; $D_2O$): δ171.16, 166.24, 151.83, 141.72, 102.67, 94.62, 88.57, 83.26, 83.20, 73.78, 72.12, 69.69, 68.42, 67.58, 65.04, 61.06, 51.66, 49.96. HRMS (ESI) *m/z* calculated for $C_{17}H_{26}N_6O_{17}P_2$ (M-H) 647.0762, found 647.0762.

Uridinyldiphosphate-α-2-deoxy-2-(4-pentynoyl)amido-D-galactopyranoside (UDP-GalNAlk; **7**):

For a 10 mg scale (39 µmol) reaction, UDP-GalNAlk (**7**) was obtained as a white powder (6.5 mg; 26% isolated yield). Product was stored at -80 °C. Characterization data are consistent with that previously reported.[12] $^1$H NMR (600 MHz; $D_2O$): δ 7.96 (d, *J* = 8.1 Hz, 1H), 5.98 (m, 2H), 5.55 (dd, *J* = 7.0, 3.5 Hz, 1H), 4.39-4.34 (m, 2H), 4.31-4.18 (m, 5H), 4.05 (d, *J* = 3.2 Hz, 1H ), 3.97 (dd, *J* = 11, 3.2 Hz, 1H), 3.78-3.73 (m, 2H), 2.60-2.49 (m, 4H), 2.36 (t, *J* = 2.6 Hz, 1H). $^{13}$C NMR (150 MHz; $D_2O$): δ175.38, 166.26, 151.83, 141.67, 102.66, 94.84, 88.52, 83.71, 83.23, 83.16, 73.78, 72.04, 69.98, 69.64, 68.42, 67.61, 65.01, 64.97, 61.01, 59.53, 49.74, 34.32, 14.32. HRMS (ESI) *m/z* calculated for $C_{20}H_{29}N_3O_{17}P_2$ (M-H) 644.0904, found 644.0899.

Uridinyldiphosphate-α-2-deoxy-2-(4,4'-azo-pentanoyl)amido-D-galactopyranoside (UDP-GalNDAz; **8**):

For a 10 mg (35 µmol) scale reaction, UDP-GalNDAz (**8**) was obtained as a white powder (4.7 mg; 20% isolated yield). Product was stored at -80 °C. Characterization data are consistent with that previously reported.[12] $^1$H NMR (600 MHz; $D_2O$): δ 7.99 (d, *J* = 8.2 Hz, 1H), 6.00 (m, 2H), 5.53 (dd, *J* = 7.0, 3.5 Hz, 1H), 4.42-4.35 (m, 2H), 4.32-4.24 (m, 3H), 4.23-4.18 (m, 2H), 4.06 (d, *J* = 3.2 Hz, 1H), 3.98 (dd, *J* = 11.0, 3.2 Hz, 1H), 3.93 (d, *J* = 5.7 Hz, 1H), 3.78-3.75 (m, 2H), 2.38-2.24 (m, 2H), 1.70 (td, *J* = 7.6, 1.1 Hz, 2H), 1.04 (s, 3H). $^{13}$C NMR (150 MHz; $D_2O$): δ 176.13, 166.24, 151.82, 141.73, 102.65, 94.78, 88.57, 83.23, 83.17, 73.78, 72.04, 69.65, 68.44, 67.64, 65.02, 62.48, 61.01, 49.76, 49.70, 30.17, 29.81, 26.32, 18.52. HRMS (ESI) *m/z* calculated for $C_{20}H_{31}N_5O_{17}P_2$ (M-H) 674.1122, found 674.1128.

4-methylumbelliferyl-*N*-acetyl-β-D-lactosaminide (LacNAcβ-MU; **9**)

GlcNAcβ-MU (10 mg, 26 mmol) and UDP-Gal (15 mg, 32 mmol.) were dissolved in a solution containing HEPES (50 mM, pH 7.5), $MnCl_2$ (2 mM), and $CaCl_2$ (1 mM). To this solution, the enzymes B4GalT1 (10 mg/mmol substrate) and mAP (10 mg/mmol substrate) were added. The reaction mixture was incubated overnight at 37 °C with gentle shaking. Reaction progress was monitored by TLC analysis (7:3 EtOH/1M $NH_4HCO_3$). Enzymes were removed by centrifugation using an Amicon® Ultra-30 centrifugal filter (10 kDa MWCO). The resulting filtrate was lyophilized and desalted on a P-2 Bio-Gel® column with 50 mM $NH_4HCO_3$ as the elution buffer to afford the product of interest. LacNAcβ-MU (**9**) was obtained as a mixture with GlcNAcβ-MU (8 mg, mixture of 91% LacNAcβ-MU and 9% GlcNAcβ-MU ) as a white powder and was stored at room temperature. Characterization data are consistent with that previously reported.[14] $^1$H NMR (600 MHz; $D_2O$): δ 7.69 (d, *J* = 8.8 Hz, 1H), 7.05 (dd, *J* = 2.4, 8.8 Hz, 1H), 7.01 (d, *J* = 2.5 Hz, 1H), 6.24 (s, 1H), 5.31 (d, *J* = 8.4 Hz, 1H), 4.55 (d, *J* = 7.8 Hz, 1H), 4.11 (t, *J* = 9.2 Hz, 1H), 4.07 (d, *J* = 11.3 Hz, 1H), 3.96 (d, *J* = 3.3 Hz, 1H), 3.93 (dd, *J* = 4.3, 12.3 Hz, 1H), 3.90-386 (m, 3H), 3.83-3.75 (m, 3H), 3.71 (dd, *J* = 3.3, 10.0 Hz, 1H), 3.60 (m, 1H), 2.43 (s, 3H), 2.08 (s, 3H). $^{13}$C NMR (150 MHz; $D_2O$): δ 174.9, 164.6, 159.7, 156.2, 154.0, 126.8, 114.0, 111.5, 103.7, 103.0, 98.8, 78.2, 75.4, 75.2, 72.6, 72.1, 71.1, 68.6, 62.5, 61.1, 59.9, 55.0, 22.2, 18.0. MS (ESI) *m/z* calculated for $C_{24}H_{31}NO_{13}$ (M+$NH_4$) 559.21, found 559.20.

General enzymatic synthesis of trisaccharide derivatives (GlcNAcR-β1,3-LacNAcβ-MU; **10-13**):

UDP-GlcNAc derivative **1-4** (0.5 mg, 0.9 mmol) was dissolved in a HEPES buffered solution (50 mM, pH 7.5) containing LacNAcβ-MU **9** (0.9 mmol), $MnCl_2$ (2 mM), and $CaCl_2$ (1 mM). To this solution, B3GNT2 (10 mg/mmol substrate) and mAP (10 mg/mmol substrate) were added. The reaction mixture was incubated overnight at 37 °C with gentle shaking. The enzymes were removed by centrifugation using an Amicon® Ultra-30 centrifugal filter (10 kDa MWCO). The resulting filtrate was loaded onto a Sep-Pak® C18 1cc Vac solid-phase extraction Cartridge (Waters). Reaction byproducts were eluted first with two column volumes of water, and the product was then eluted with 50% acetonitrile. The extracted trisaccharide products (**10-13**) were lyophilized and analyzed by HPLC-MS (Figure 2B,D).

General enzymatic synthesis of tetrasaccharides derivatives (LacNR-β1,3-LacNAcβ-MU; **14-17**):

Trisaccharide derivatives **10-13** were dissolved in a HEPES buffered solution (50 mM, pH 7.5) containing UDP-Gal (0.9 mmol, 1 equiv.), $MnCl_2$ (2 mM), and $CaCl_2$ (1 mM). To this solution, B4GalT1 (10 mg/mmol substrate) and mAP (10 mg/mmol substrate) were added. The reaction mixture was incubated overnight at 37 °C with gentle shaking. The enzymes were removed by centrifugation using an Amicon® Ultra-30 centrifugal filter (10 kDa MWCO). The resulting filtrate was loaded onto a Sep-Pak® C18 1cc Vac solid-phase extraction Cartridge (Waters). Reaction byproducts were eluted first with two column volumes of water, and the product was then eluted with 50% acetonitrile. The extracted tetrasaccharide products (**14-17**) were lyophilized and analyzed by HPLC-MS (Figure 2C,D).

**HPLC-MS Purification Conditions of Nucleotide Sugars**

All nucleotide-sugar derivatives (**1-8**) were purified by HPLC using a Waters© XBridge® BEH Amide column (130Å, 5 mm, 10 x 250 mm, 1/pkg) with 1% of the flow diverted to the ESI-MS detector. UV absorbance was monitored at 264 nm and the system was controlled *via* Agilent© OpenLab ChemStation chromatography software. Mobile phase A was ammonium formate in water (10 mM, adjusted to pH 4.5 with formic acid); Mobile phase B was a mixture of acetonitrile (90%) with ammonium formate in water (10%; 10 mM, pH = 4.5 with formic acid). The following gradient was used with a flow rate of 5 mL/min and an injection volume of 100 µL (20 mg of compound): 1) Gradient of 85%-50% mobile phase B from 0-40 min, 2) gradient of 35%-15% mobile phase B from 40-45 min, 3) isocratic run of 15% mobile phase B from 45-50 min, 4) gradient of 15%-85% mobile phase B from 50-55 min, 5) followed by a final equilibration isocratic run of 85% mobile phase B from 55-60 min.

**HPLC-MS Analytical Conditions of Oligosaccharides**

All oligosaccharides (**10-17**) were analyzed by HPLC-MS using a Waters© XBridge® BEH Amide column (130Å, 5 mm, 4.6 x 250 mm, 1/pkg) with 1% of the flow diverted to the ESI-MS detector. UV absorbance was monitored at 315 nm and the system was controlled *via* Agilent© OpenLab ChemStation chromatography software. Mobile phase A was ammonium formate in water (10 mM, adjusted to pH 4.5 with formic acid); Mobile phase B was a mixture of acetonitrile (90%) with ammonium formate in water (10%; 10 mM, pH = 4.5 with formic acid). The following gradient was used with a flow rate of 2 mL/min and an injection volume of 5 µL (0.2 mg of compound): 1) Isocratic run of 100% mobile phase B from 0-10 min, 2) gradient of 100%-90% mobile phase B from 10-35 min, 3) gradient of 90%-50% mobile phase B from 35-45 min, 4) isocratic run of 50% mobile phase B from 45-50 min, 5) gradient of 50%-100% mobile phase B from 50-55%, followed by a final equilibration isocratic run of 100% mobile phase B from 55-60 min.

#### Biological Methods

##### Cell staining materials

AZDye 405 DBCO (DBCO-405) was purchased from Click Chemistry Tools (cat. 1310); DBCO-s-biotin was purchased from Sigma (cat. 760706); Streptavidin-PacificBlue conjugate (cat. S11222) was purchased from Thermo-Fisher Scientific; Griffonia (Bandeiraea) Simplicifolia Lectin II Biotinylated (GSL-II; cat. VECTB1215) was purchased from MJS BioLynx Inc. Revert Revert™ 700 Total Protein Stain (cat. 926-11021) and Streptavidin-IRDye® 800CW (cat. 926-32230) were purchased from Li-Cor Biosciences.

##### Cell culture

Jurkat and U937 cells were cultured in RPMI-1640 medium (with L-glutamine, sodium bicarbonate) supplemented with 10% fetal bovine serum (FBS) and 1× penicillin/streptomycin (P/S). Suspension cells were passaged after reaching $2\times 10^6$ viable cells/mL. HS-578-T and HEK293T cells were cultured in high-glucose DMEM medium with L-glutamine, supplemented with 10% fetal bovine serum (FBS) and 1× penicillin/streptomycin (P/S). Adherent cells were passaged using 0.25% trypsin-EDTA after reaching ~80% confluency. All cells were maintained in a humid 5% $CO_2$ atmosphere at 37°C.

##### Cell-surface glycan labeling

Adherent cells were plated in 12-well plates (300 000 cells/well) and grown to 80% confluency. Suspended cells were centrifuged and resuspended at 6 M cells/mL and 200 uL was aliquoted per cell treatment. Prior to labeling, cells were washed with a culture medium without FBS and P/S. Washed cells were incubated in a mixture of serum-free culture medium containing 50 µg/mL *C. perfringens* neuraminidase (NanH), 250 µg/mL B3GNT2, the designated concentration for the experiment of either UDP-GlcNAc (**1**) or UDP-GlcNAz (**2**), 0.1% BSA, and 5 mM $MnCl_2$ for 2 hours at 37°C. Untreated control experiments were treated with a mixture of serum-free culture medium containing 0.1% BSA, 5 mM $MnCl_2$, 50 µg/mL NanH with or without B3GNT2 or the appropriate sugar nucleotide (**1** or **2**). For LacNAc/LacNAz extension, the GlcNAc/GlaNAz engineered cells were washed with serum-free culture medium and further reacted in a mixture of serum-free culture medium containing 250 µg/mL B4GalT1, 0.2 mM UDP-Gal, 0.1% BSA, and 5 mM $MnCl_2$ for 2 hours at 37°C. Untreated control experiments were treated with a mixture of serum-free culture medium containing 0.1% BSA and 5 mM $MnCl_2$, with or without B4GalT1 or UDP-Gal. Following glycan labeling, cells were washed three times with 1% FBS/DPBS and then treated as indicated.

##### Flow cytometry analysis

For the detection of cell-surface glycan labeling with UDP-GlcNAz (**2**) by DBCO-405 on adherent cells, the cells were washed three times with DPBS with Ca/Mg and then treated with 300 uL staining buffer (DPBS with Ca/Mg) with 50 uM DBCO-405 and incubated at room temperature in the dark for 1 hour. Cells were then washed two times with DPBS without Ca/Mg and detached using Cell Dissociation Buffer, enzyme free (cat. 13151014) from Thermo Fisher. Cells were then suspended in 1% FBS/DPBS, centrifuged gently (300 rpm for 3 min), washed twice with DPBS with Ca/Mg, resuspended in 500 uL FACS buffer (PBS without Ca/Mg supplemented with 2 mM EDTA and 0.5% BSA) and transferred to tubes for flow cytometric analysis (Beckman Coulter, Cytoflex S).

For the detection of cell-surface glycan labeling with UDP-GlcNAz (**2**) by DBCO-405 on suspension cells, the cells were washed three times with DPBS with Ca/Mg, centrifuged gently (300 rpm for 3 min), and then treated with 100 uL staining buffer (DPBS with Ca/Mg) with 50 uM DBCO-405 and incubated at room temperature in the dark for 1 hour. Cells were then washed twice with DPBS with Ca/Mg, resuspended in 500 uL FACS buffer (PBS without Ca/Mg supplemented with 2 mM EDTA and 0.5% BSA) and transferred to tubes for flow cytometric analysis (Beckman Coulter, Cytoflex S).

For the detection of GlcNAc-, GlcNAz-, LacNAc- or LacNAz-engineered Jurkat cells, glcyo-engineered cells were washed three times with 1% FBS/DPBS, centrifuged gently (300 rpm for 3 min), and then treated in 100 uL staining buffer (1% FBS/DPBS) with 1 ug/mL GSL-II for 30 min at 4°C. Cells were then washed once with staining buffer, centrifuged and then treated in 100 uL staining buffer (1% FBS/DPBS) with 2.5 ug/mL Strep-PB for 30 min at 4°C. Cells were then washed twice with 1% FBS/DPBS, resuspended in 500 uL FACS buffer (PBS without Ca/Mg supplemented with 2 mM EDTA and 0.5% BSA) and transferred to tubes for flow cytometric analysis (Beckman Coulter, Cytoflex S).

For all treatments, cell viability was determined by adding 1 ug/mL PI to cell suspensions 1 min prior to analysis. The live population cells were gated based on forward and side scatter emission and exclusion of PI positive cells on the PE-A (585/42 BP filter) emission channel. Strep-PB and DBCO-405 binding were both determined by fluorescence intensity on the PB450 (405/45 BP filter).

**Immunoblot analysis**

After cell surface glycan labeling of adherent cells with UDP-GlcNAz (**2**), cells were lysed with RIPA lysis buffer containing 1× protease inhibitor cocktail (NEB cat # 5871S). Cell lysates were clarified by centrifugation at 14 000*g* for 15 min and the total protein content of supernatants was assessed by BCA assay (Pierce, Thermo Fisher). Samples (15 ug total protein for HEK293T cells and 20 ug total protein for HS-578-T cells) were resolved on an 8% SDS-PAGE gel and transferred to a low-fluorescent PVDF membrane (Immobilon-FL, Sigma). For normalization, total protein was stained with Revert Total Protein stain (Li-Cor Biosciences) for 5 min, washed twice with wash buffer (6.7% acetic acid, 30% methanol) and rinsed briefly in TBS prior to scanning on an Odyssey Li-Cor CLx scanner (Li-Cor Biosciences). Next, the membrane was blocked in a blocking buffer (5% nonfat dry milk in TBS) for 1 hour at room temperature. The blocked membrane was incubated for 1 hour at room temperature with Strep-800 (1:1000) in blocking buffer with 0.1% Tween-20 (TBST) and washed with TBST (3 × 5 min). Membranes were rinsed briefly in TBS prior to imaging. Final detection of Strep-800 was performed on an Odyssey Li-Cor CLx scanner (Li-Cor Biosciences).

**Enzyme Expression and Purification**

Bacterial proteins *P. multocida N*-acetylglucosamine-1-phosphate wild-type and mutant uridylyltransferases (PmGlmU and PmGlmU$^{T199A}$), *P. multocida* strain Pm70 inorganic pyrophosphatase (PmPpA), *B. longum* strain ATCC55813 *N*-acetylhexosamine-1-kinase (NahK) *C. perfringens* α2,3/6/8 sialidase (NanH), *S. pneumoniae* β-galactosidase (BgaA), metagenomic alkaline phosphatase (mAP), and the human UDP-N-acetylhexosamine wild-type and mutant pyrophosphorylases (AGX1 and AGX1$^{F383A}$) were recombinantly expressed as previously reported (Table S1). The genes encoding PmGlmU, PmGlmU$^{T199A}$, AGX1, AGX1$^{F383A}$, and NanH (GenBank: Y00963.1) were commercially synthesized and inserted into pET-15b vectors using the

NdeI and XhoI restriction sites (Genscript). The gene encoding PmPpA was commercially synthesized and inserted into a pET-15b vector using NdeI and BamHI restriction sites (Genscript). The gene encoding metagenomic alkaline phosphatase (mAP) was commercially synthesized and ligated into a pET-21a(+) plasmid using NdeI and XhoI restriction sites (Genscript). The gene encoding NahK was commercially synthesized and inserted into a pET-22b(+) vector using NdeI and XhoI restriction sites (Genscript).

General procedure for bacterial enzyme expression and purification

See **Table S1** for specific expression details for all bacterial enzymes used in this work. *E. coli* BL-21 cells transformed with plasmid containing genes encoding enzyme of interest and ampicillin resistance were cultured in 500 mL or 1 L LB-Miller medium with ampicillin (100 μg/mL) at 37°C with gentle shaking (200 rpm) until the desired $OD_{600nm}$ was reached. Protein expression was induced by the addition of 0.1mM of isopropyl-1-thio-β-D-galactopyranoside (IPTG) and cultures were incubated at a specified temperature with rigorous shaking (250 rpm) for a specified number of hours. The cells were harvested by centrifugation (4,000 rpm) at 4°C for 1 h and the resulting pellet was resuspended in lysis buffer (100 mM Tris-HCl, 0.1% Triton X-100, pH 8.0, 100 mg/mL lysozyme, and 5 mg/mL protease inhibitor; 5 mL/g cell pellet). The cells were then lysed by passing the suspension twice through an Emulsiflex high pressure homogenizer. Subsequently, the lysate was centrifuged (12,000 rpm) at 4°C for 45 minutes to obtain a cell pellet. Enzyme ($His_6$-tagged) purification was performed by loading the lysis supernatant onto a Ni-NTA column pre-equilibrated with binding buffer (10 mM imidazole, 500 mM NaCl, 50 mM Tris-HCl, pH 7.5). The loaded column was then washed with binding buffer (10 mM imidazole, 500 mM NaCl, 50 mM Tris-HCl, pH 7.5) followed by wash buffer (50 mM imidazole, 500 mM NaCl, 50 mM Tris-HCl, pH 7.5). The enzyme of interested was eluted with elution buffer (200 mM imidazole, 500 mM NaCl, 50 mM Tris-HCl, pH 7.5) followed by a high concentration elution buffer (500 mM imidazole, 500 mM NaCl, 50 mM Tris-HCl, pH 7.5). Isolation of the enzyme was confirmed by SDS-PAGE and the fractions containing purified enzyme were combined, buffer exchanged as indicated in Table S1 and concentrated by centrifugation (4,000 rpm at 4 °C) using an Amicon® Ultra-30 centrifugal filter (10 kDa MWCO). Enzyme concentration was determined by BCA protein assay (Pierce, Thermo Fisher), and the enzyme was kept in 20% glycerol for long-term storage at a specified temperature indicated in Table S1.

**Table S1**. Specifications and deviations from general bacterial expression & purification.

| Enzyme | Species | Activity | Vector | OD | Induction Temp. & Time | Buffer Exchange Method | Storage Conditions | Yield (mg/500 mL) | Ref. |
|---|---|---|---|---|---|---|---|---|---|
| PmGlmU | *Pasteurella multocida* | *N*-acetyl-glucosamine 1-phosphate uridylyl-transferase | pET-15b | 0.8-1.0 | 25°C for 18 h | dialysis | 25 mM Tris-HCl, pH 7.5, -20°C | 36 | 11 |
| PmGlmU$^{T199A}$ | *Pasteurella multocida* | *N*-acetyl-glucosamine 1-phosphate uridylyl-transferase | pET-15b | 0.8-1.0 | 25°C for 18 h | dialysis | 25 mM Tris-HCl, pH 7.5, -20°C | 23 | - |
| AGX1 | Human | UDP-*N*-acetyl-hexosamine pyro-phosphorylase | pET-15b | 0.6-0.8 | 16°C for 20 h | dialysis | 50 mM Tris-HCl, pH 7.5, -20°C | 14 | 13 |
| AGX1$^{F383A}$ | Human | UDP-*N*-acetyl-hexosamine pyro-phosphorylase | pET-15b | 0.6-0.8 | 16°C for 20 h | dialysis | 50 mM Tris-HCl, pH 7.5, -20°C | 12 | - |
| NahK | *Bifido-bacterium Longum* strain ATCC55813 | N-acetyl-hexosamine 1-kinase | pET-22b(+) | 0.8-1.0 | 20°C for 24 h | Amicon® Ultra-30 centri-fugation filter | 20 mM Tris-HCl, pH 7.5, -80°C | 26 | 15 |
| PmPpA | *Pasteurella multocida* strain Pm70 | Inorganic pyrophosphatase | pET-15b | 0.8-1.0 | 25°C for 20 h | Amicon® Ultra-30 centri-fugation filter | 20 mM Tris-HCl, pH 7.5, -80°C | 140 | 14 |
| NanH | *Clos-tridium per-fringens* | *a2-3,-6,-8* Neuraminidase | pET-15b | 0.6-0.8 | 25°C for 16 h | Amicon® Ultra-30 centri-fugation filter | 20 mM Tris-HCl, pH 7.5, -80°C | 127 | 16 |
| BgaA | *Strepto-coccus pneu-moniae* | *β1,4* Galactosidase | pET-28b(+) | 0.6-0.8 | 20°C for 18 h | Amicon® Ultra-30 centri-fugation filter | 25mM Tris-HCl, , 0.1 M NaCl, pH 7.5, -80°C | 151 | 17 |
| mAP | Meta-genomic | Alkaline phosphatase | pET-21a(+) | 0.4-0.5 | 15°C for 40 h | Amicon® Ultra-30 centri-fugation filter | 20 mM Tris-HCl, pH 7.5, -80°C | 19 | 18 |

General procedure for human glycosyltransferase expression and purification

The plasmids encoding soluble, secreted GFP fusion proteins containing the catalytic domain of human β1,3-*N*-acetylglucosaminlytransferase 2 (B3GNT2) or β1,4-galactosyltransferase 1 (B4GalT1) in pGEn2-DEST vectors were purchased from DNASU (cat. HsCD00413085, HsCD00522296). Recombinant GFP-B3GNT2 and GFP-B4GalT1 protein was expressed and purified as previously described using Expi293 cells.[16, 19] Plasmids were chemically transformed into DH5-alpha competent cells. Transformed cells were cultured in 5 mL LB media supplemented with 100 μg/mL ampicillin for 16 h at 37 °C and shaking (200 rpm) for 16 hours. Cultured cells were then scaled up to 500 mL and plasmids were isolated using the PureLink™ HiPure Plasmid Maxiprep Kit (Thermofisher, cat. K210006).

Expi293 cell maintenance

Expi293 cells (ThermoFisher) were cultured in Expi293 Expression Medium (ThermoFisher). Cells were maintained in a humid 5% CO2 atmosphere at 37°C with shaking at 120 rpm. Cells were passaged for maintenance after reaching $4 \times 10^6$ viable cells/mL. Cells were cultured for at least 3 passages following thaw prior to transfection.

Transfection and Expression

Expi293 cells were transiently transfected with the vector containing the B3GNT2 or B4GalT1 gene using the Expifectamine 293 Transfection Kit (ThermoFisher, cat. A14524). On day 5 following transfection, cells were harvested by centrifugation (25 min, 4000 rcf). The supernatant was collected and adjusted to contain 20 mM imidazole, 200 mM NaCl, and 30 mM sodium phosphate, at pH 7.2. The resulting solutions were then filtered using a 0.45 μm PES Filter Unit and applied to a Ni2+-sepharose resin pre-equilibrated with column buffer (20 mM HEPES, 300 mM NaCl, pH 7.2) containing 20 mM imidazole. The column was sequentially washed with column buffer containing 20 mM, 50 mM, and 100 mM imidazole prior to elution with 300 mM imidazole, and the column was flushed with 500 mM. Overexpression and purification were assessed by SDS-PAGE. Purified protein was buffer exchanged into 20 mM Tris-HCl (pH 7.5) buffer, concentrated using a 10 kDa MWCO spin filter (Amicon), and stored in 20 mM Tris-HCl (pH 7.5) with 20% glycerol at -80°C. The final protein concentration was determined by BCA assay. Yields per 100 mL culture were: B3GNT2 = 8.7 mg; B4GalT1 = 4.0 mg.

#### References

(1) Bond, M. R.; Zhang, H.; Vu, P. D.; Kohler, J. J. Photocrosslinking of glycoconjugates using metabolically incorporated diazirine-containing sugars. *Nat. Protoc.,* **2009**, *4,* 1044-1063.
(2) Gilormini, P. A.; Lion, C.; Vicogne, D.; Levade, T.; Potelle, S.; Mariller, C.; Guérardel, Y.; Biot, C.; Foulquier, F. A sequential bioorthogonal dual strategy: ManNAl and SiaNAl as distinct tools to unravel sialic acid metabolic pathways. *Chem. Commun.,* **2016**, *52,* 2318-2321.
(3) Bannwarth, W.; Knorr, R. Formation of carboxamides with N,N,N′ ,N′ -tetramethyl (succinimido) uronium tetrafluoroborate in aqueous / organic solvent systems. *Tetrahedron Lett.,* **1991**, *32,* 1157-1160.
(4) Brown, K.; Pompeo, F.; Dixon, S.; Mengin-Lecreulx, D.; Cambillau, C.; Bourne, Y. Crystal structure of the bifunctional N-acetylglucosamine 1-phosphate uridyltransferase from Escherichia coli: a paradigm for the related pyrophosphorylase superfamily. *EMBO J.,* **1999**, *18,* 4096-4107.
(5) Peneff, C.; Ferrari, P.; Charrier, V.; Taburet, Y.; Monnier, C.; Zamboni, V.; Winter, J.; Harnois, M.; Fassy, F.; Bourne, Y. Crystal structures of two human pyrophosphorylase isoforms in complexes with UDPGlc(Gal)NAc: role of the alternatively spliced insert in the enzyme oligomeric assembly and active site architecture. *EMBO J.,* **2001**, *20,* 6191-6202.
(6) Slater, M.; Snauko, M.; Svec, F.; Fréchet, J. M. J. “Click Chemistry” in the Preparation of Porous Polymer-Based Particulate Stationary Phases for μ-HPLC Separation of Peptides and Proteins. *Anal. Chem.,* **2006**, *78,* 4969-4975.
(7) Brabez, N.; Lynch, R. M.; Xu, L.; Gillies, R. J.; Chassaing, G.; Lavielle, S.; Hruby, V. J. Design, synthesis, and biological studies of efficient multivalent melanotropin ligands: tools toward melanoma diagnosis and treatment. *J. Med. Chem.,* **2011**, *54,* 7375-7384.
(8) Soares da Costa, D.; Sousa, J. C.; S, D. M.; Petkova-Yankova, N. I.; Marques, F.; Reis, R. L.; Sousa, N.; Pashkuleva, I. Bioorthogonal Labeling Reveals Different Expression of Glycans in Mouse Hippocampal Neuron Cultures during Their Development. *Molecules,* **2020**, *25,* 795.
(9) Murali, M.; Murali, V. P.; Joseph, M. M.; Rajan, S.; Maiti, K. K. Elucidating cell surface glycan imbalance through SERS guided metabolic glycan labelling: An appraisal of metastatic potential in cancer cells. *J. Photochem. Photobiol., B,* **2022**, *234,* 112506.
(10) Tanaka, Y.; Kohler, J. J. Photoactivatable Crosslinking Sugars for Capturing Glycoprotein Interactions. *J. Am. Chem. Soc.,* **2008**, *130,* 3278-3279.
(11) Chen, Y.; Thon, V.; Li, Y.; Yu, H.; Ding, L.; Lau, K.; Qu, J.; Hie, L.; Chen, X. One-pot three-enzyme synthesis of UDP-GlcNAc derivatives. *Chem. Commun.,* **2011**, *47,* 10815-10817.
(12) Wen, L.; Gadi, M. R.; Zheng, Y.; Gibbons, C.; Kondengaden, S. M.; Zhang, J.; Wang, P. G. Chemoenzymatic Synthesis of Unnatural Nucleotide Sugars for Enzymatic Bioorthogonal Labeling. *ACS Catal.,* **2018**, *8,* 7659-7666.
(13) Guan, W.; Cai, L.; Wang, P. G. Highly Efficient Synthesis of UDP-GalNAc/GlcNAc Analogues with Promiscuous Recombinant Human UDP-GalNAc Pyrophosphorylase AGX1. *Chem. Eur. J.,* **2010**, *16,* 13343-13345.
(14) Lau, K.; Thon, V.; Yu, H.; Ding, L.; Chen, Y.; Muthana, M. M.; Wong, D.; Huang, R.; Chen, X. Highly efficient chemoenzymatic synthesis of β1–4-linked galactosides with promiscuous bacterial β1–4-galactosyltransferases. *Chem. Commun.,* **2010**, *46,* 6066-6068.
(15) Li, Y.; Yu, H.; Chen, Y.; Lau, K.; Cai, L.; Cao, H.; Tiwari, V. K.; Qu, J.; Thon, V.; Wang, P. G.; et al. Substrate Promiscuity of N-Acetylhexosamine 1-Kinases. *Molecules,* **2011**, *16,* 6396-6407.

(16) Babulic, J. L.; Capicciotti, C. J. Exo-Enzymatic Cell-Surface Glycan Labeling for Capturing Glycan–Protein Interactions through Photo-Cross-Linking. *Bioconjug. Chem.,* **2022**, *33,* 773-780.
(17) Zhang, X.; Chen, F.; Petrella, A.; Chacón-Huete, F.; Covone, J.; Tsai, T.-W.; Yu, C.-C.; Forgione, P.; Kwan, D. H. A High-Throughput Glycosyltransferase Inhibition Assay for Identifying Molecules Targeting Fucosylation in Cancer Cell-Surface Modification. *ACS Chem. Biol.,* **2019**, *14,* 715-724.
(18) Lee, D.-H.; Choi, S.-L.; Rha, E.; Kim, S. J.; Yeom, S.-J.; Moon, J.-H.; Lee, S.-G. A novel psychrophilic alkaline phosphatase from the metagenome of tidal flat sediments. *BMC Biotechnol.,* **2015**, *15,* 1.
(19) Moremen, K. W.; Ramiah, A.; Stuart, M.; Steel, J.; Meng, L.; Forouhar, F.; Moniz, H. A.; Gahlay, G.; Gao, Z.; Chapla, D. Expression system for structural and functional studies of human glycosylation enzymes. *Nat. Chem. Biol.,* **2018**, *14,* 156-162.

#### NMR Characterization

[1]H and [13]C NMR Spectra of UDP-GlcNAc (**1**)

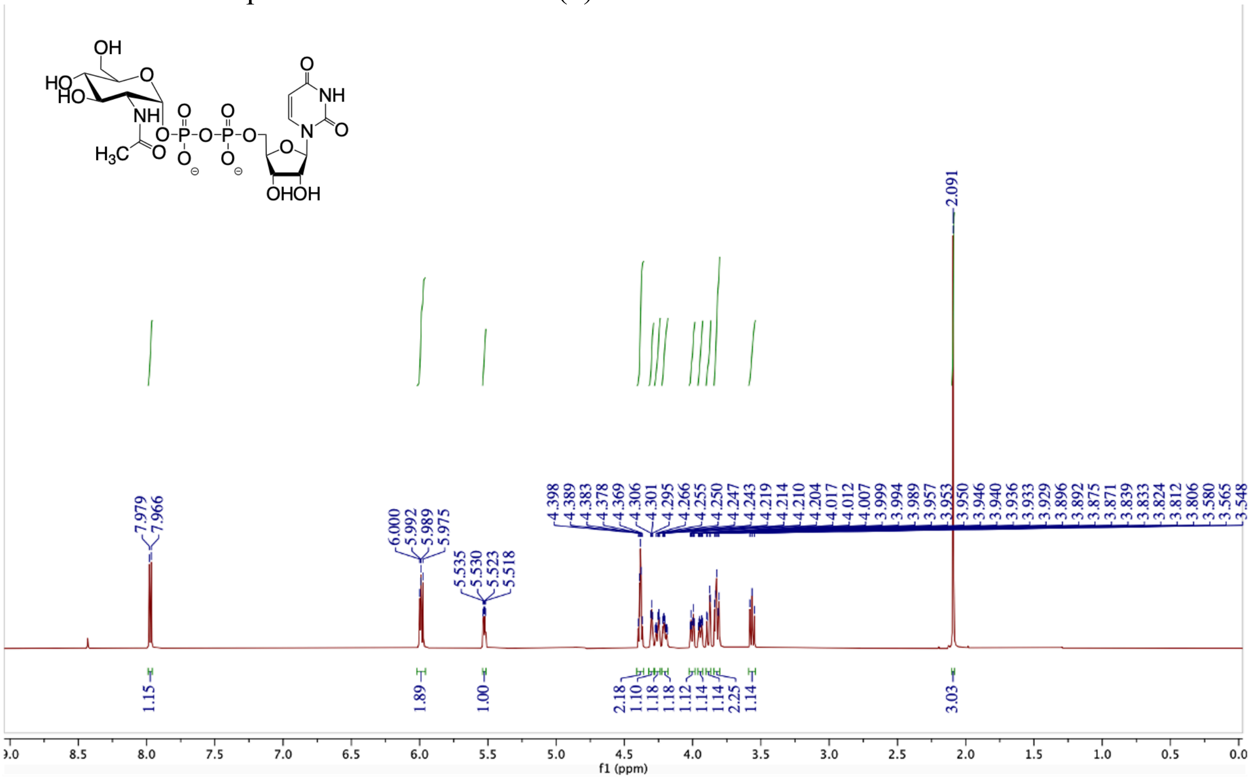

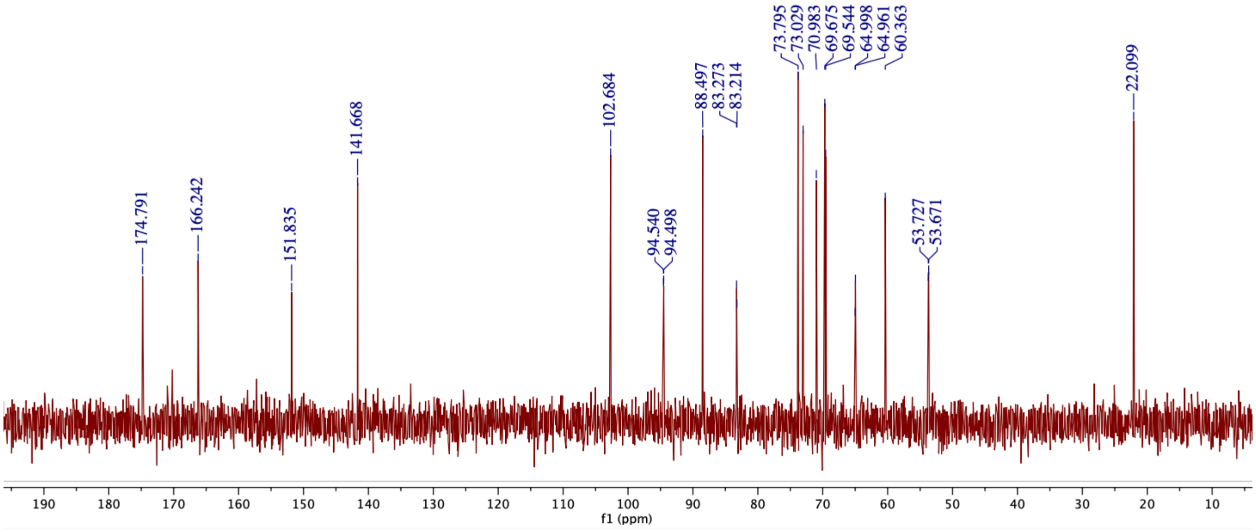

[1]H and [13]C NMR Spectra of UDP-GlcNAz (**2**)

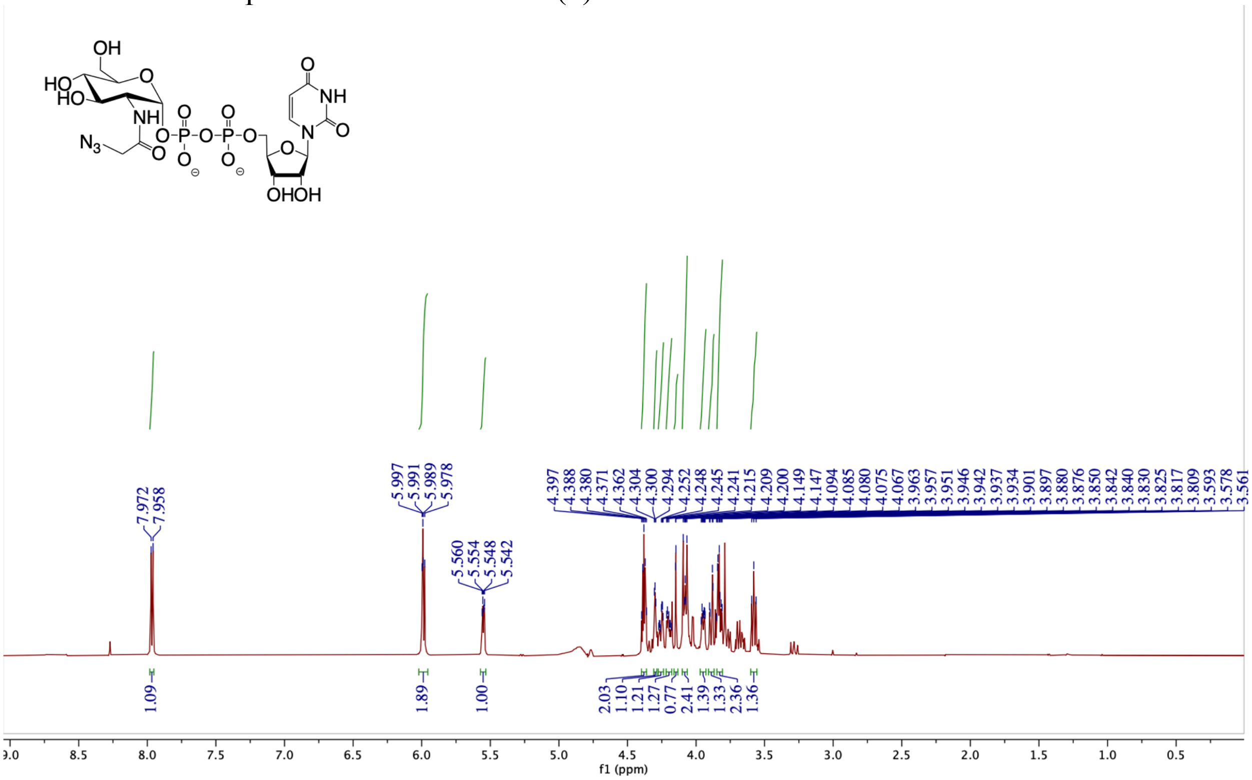


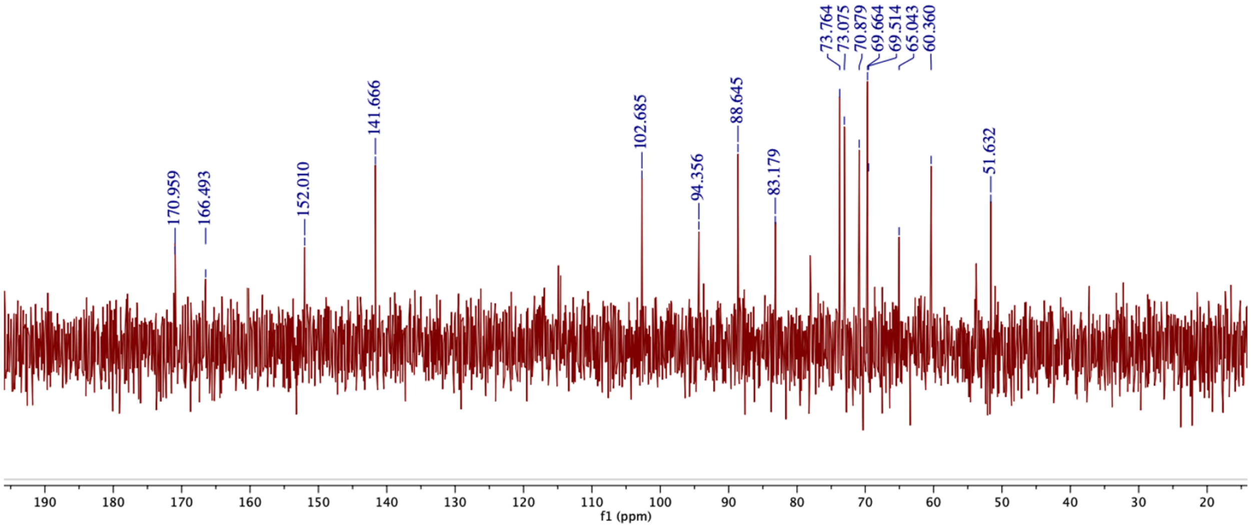

[1]H and [13]C NMR Spectra of UDP-GlcNAlk (**3**)

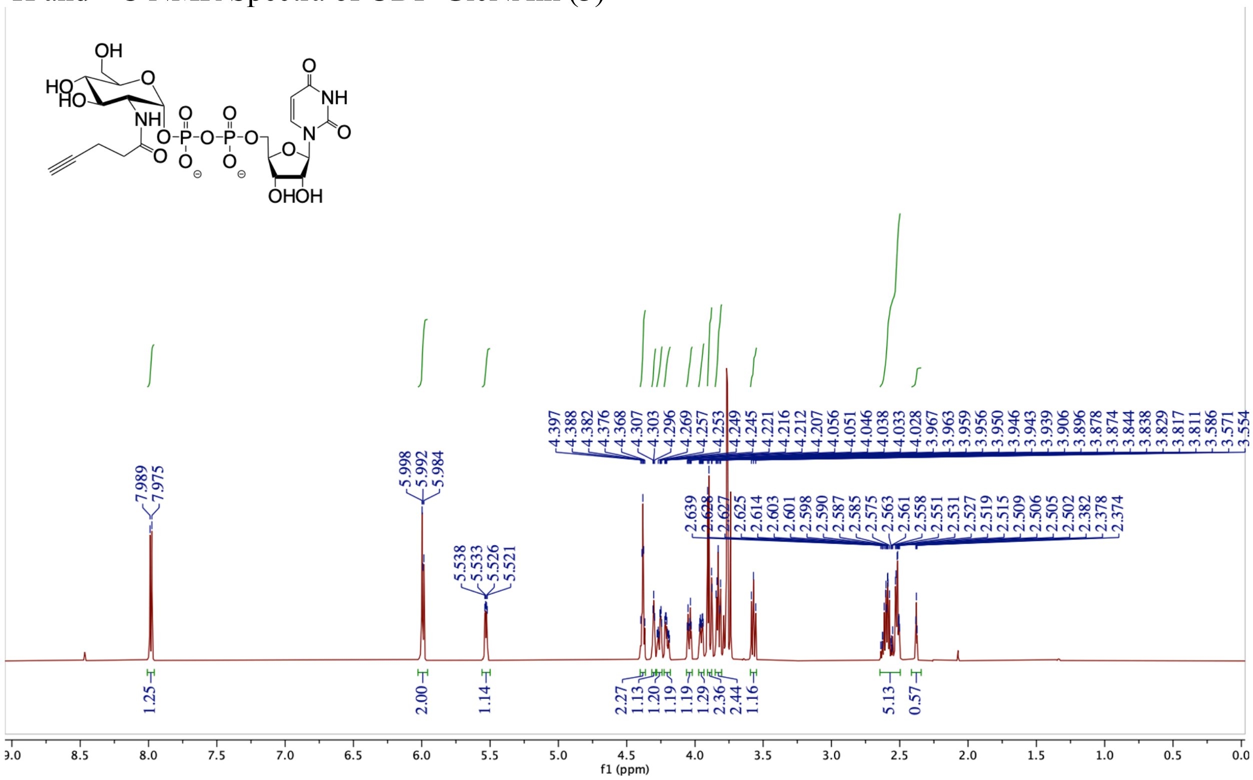


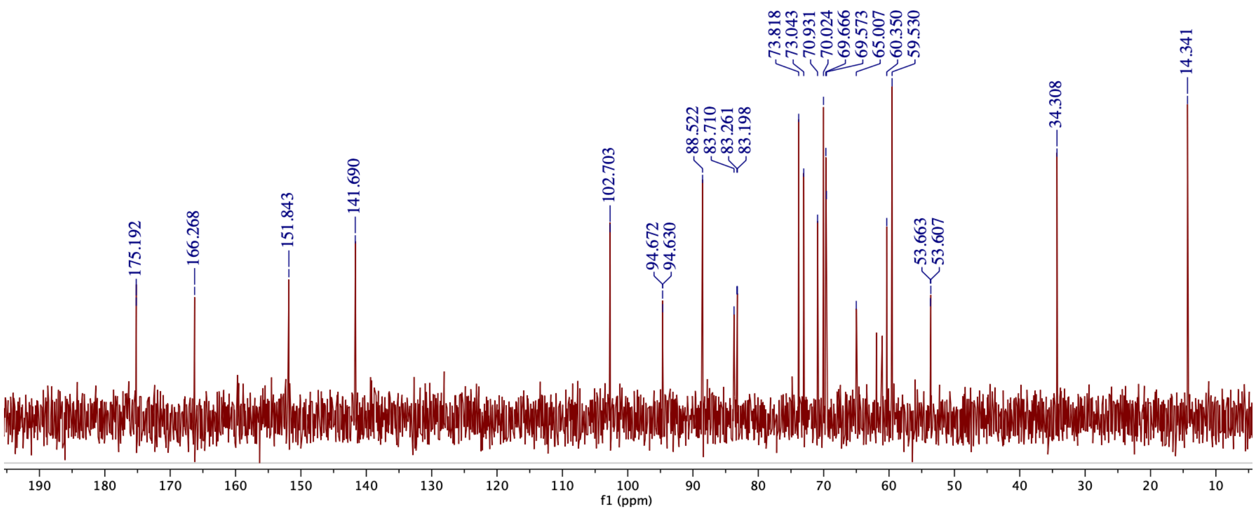

[1]H and [13]C NMR Spectra of UDP-GlcNDAz (**5**)

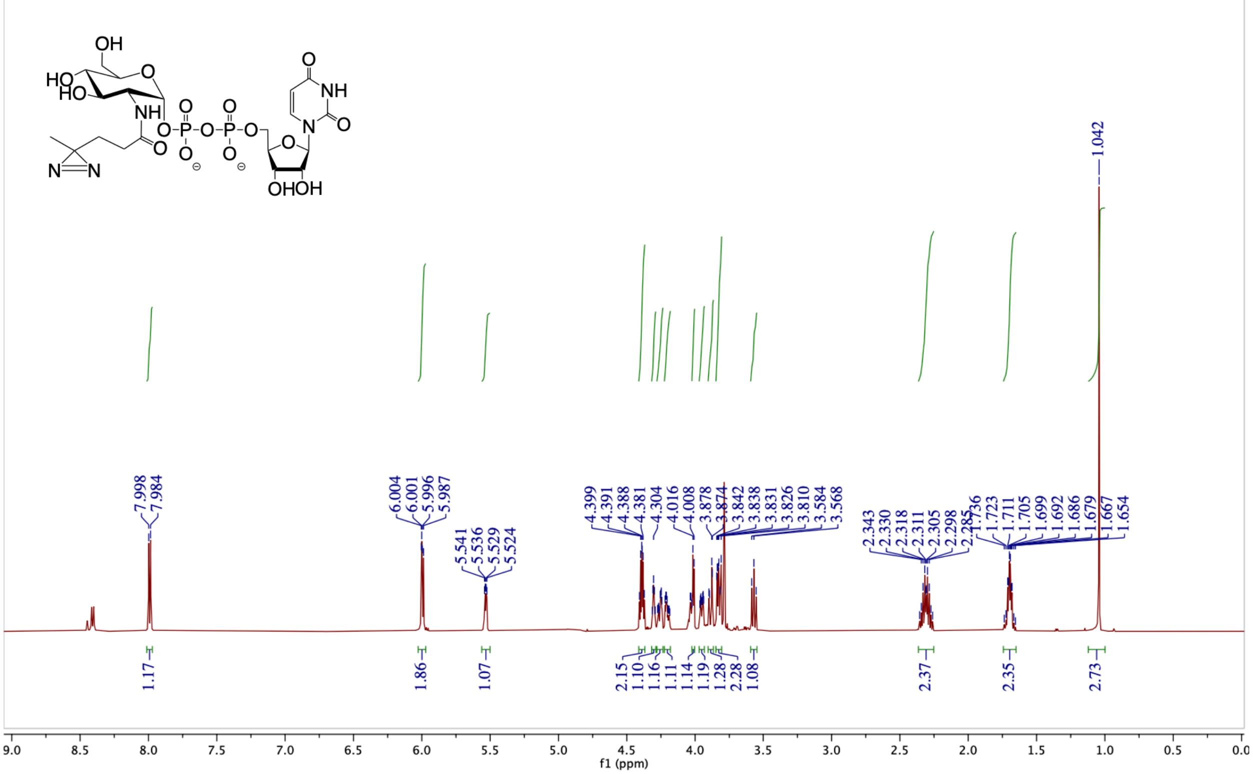


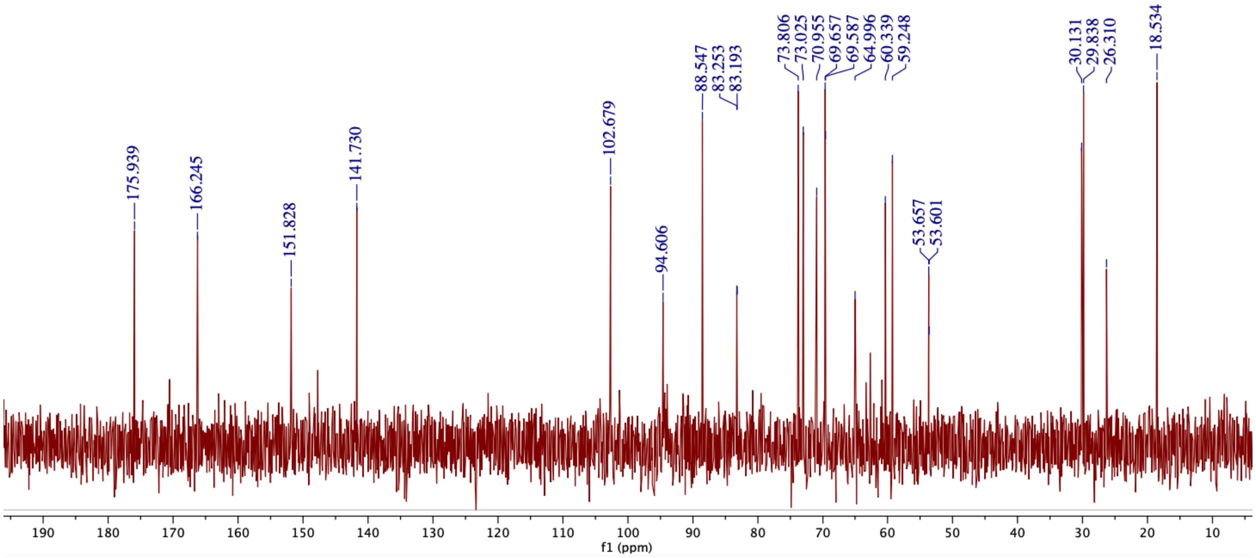

$^{1}H$ and $^{13}C$ NMR Spectra of UDP-GalNAc (**5**)

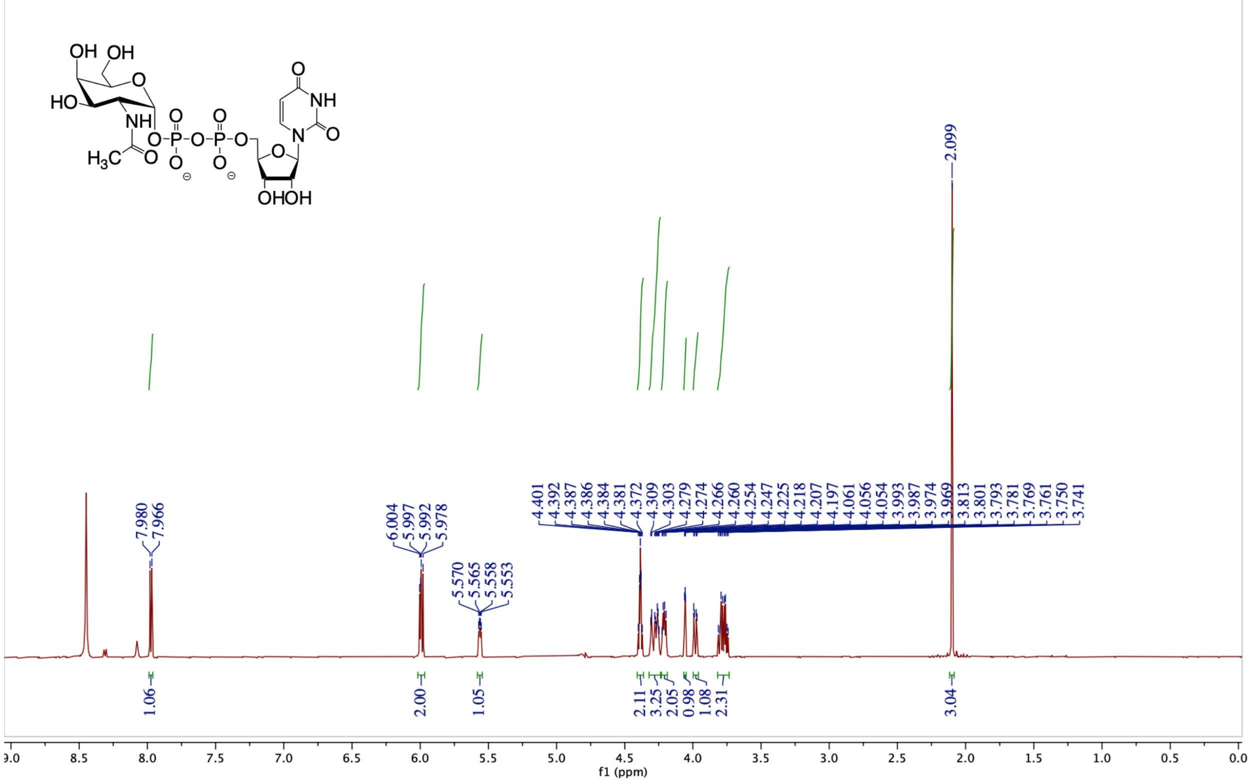

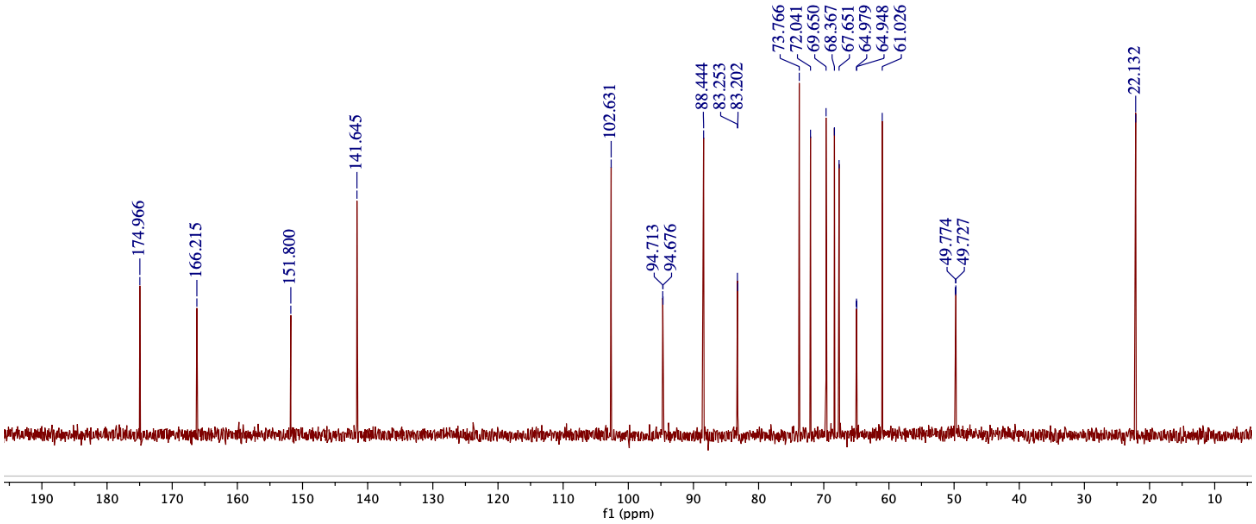

[1]H and [13]C NMR Spectra of UDP-GalNAz (**6**)

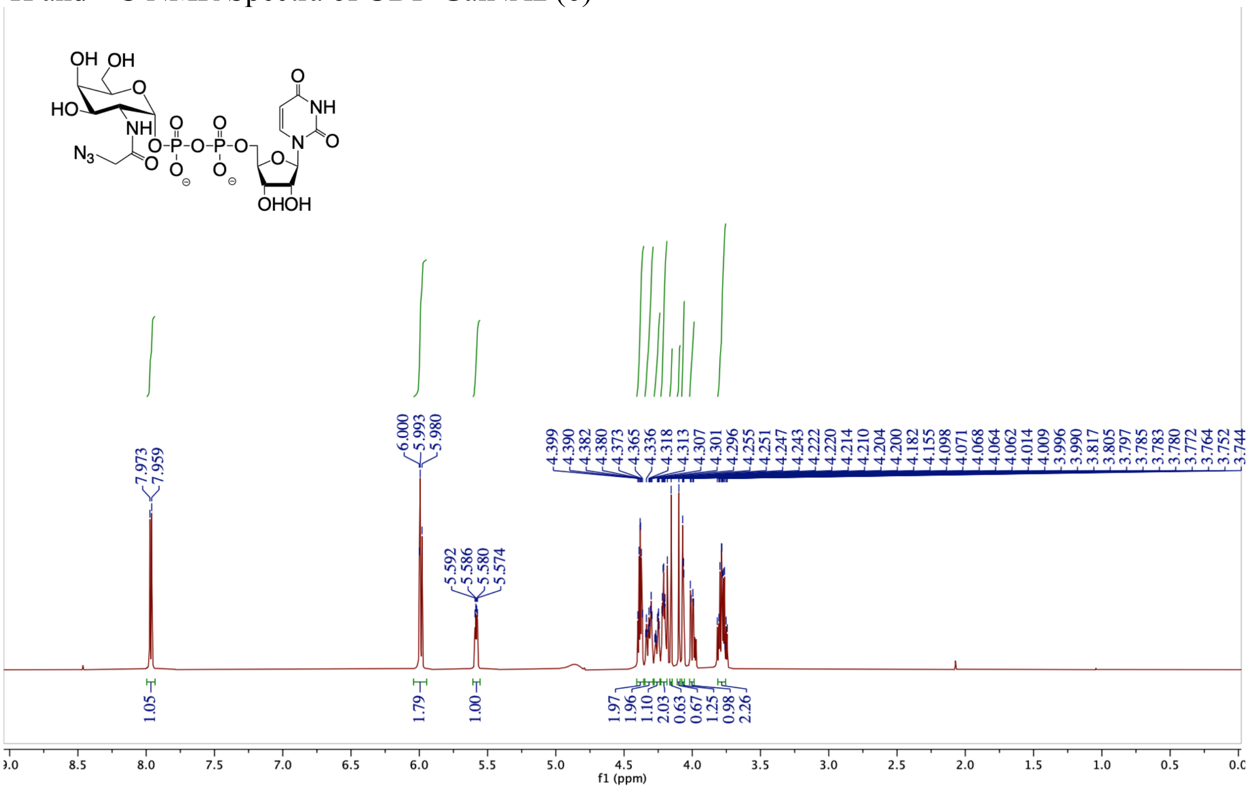


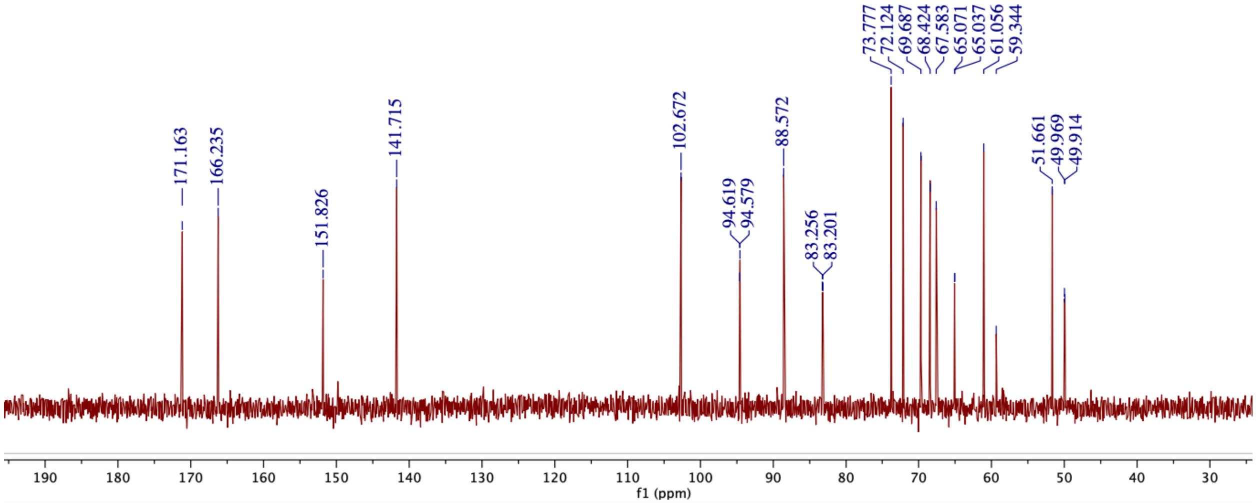

[1]H and [13]C NMR Spectra of UDP-GalNAlk (**7**)

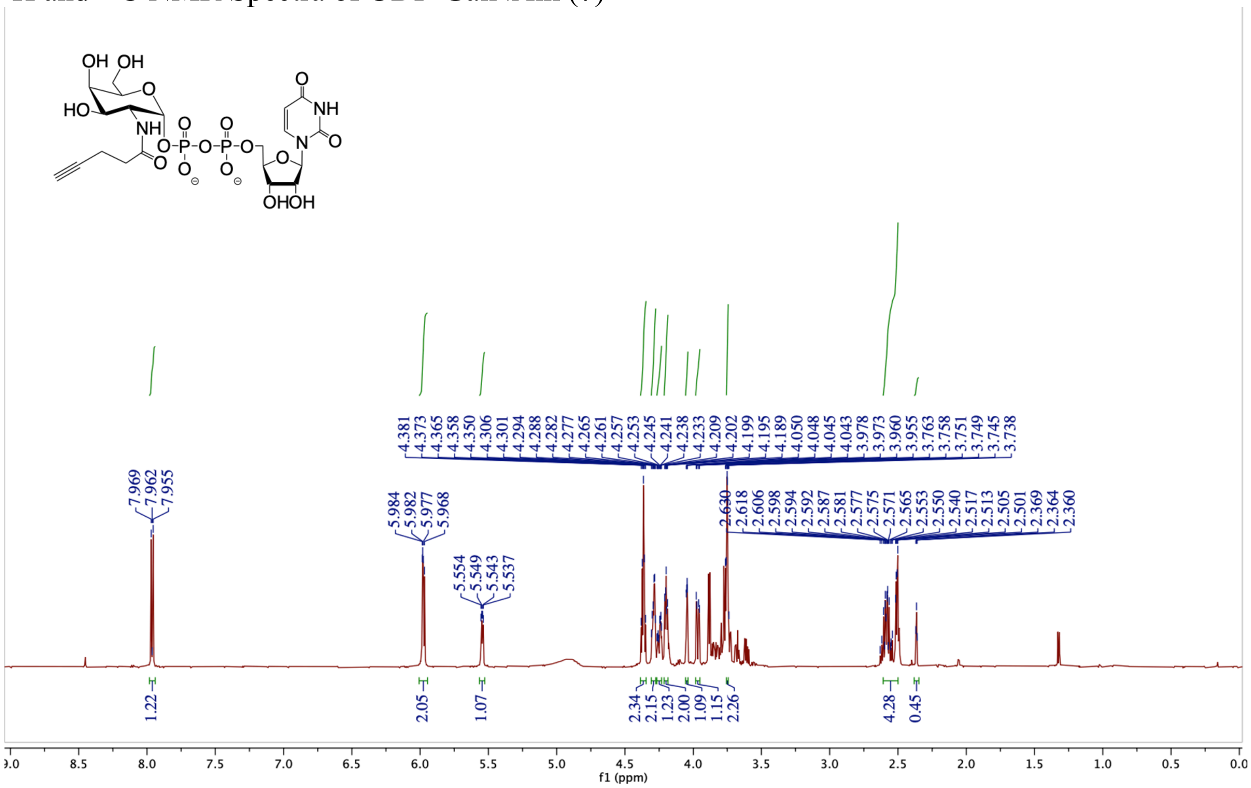


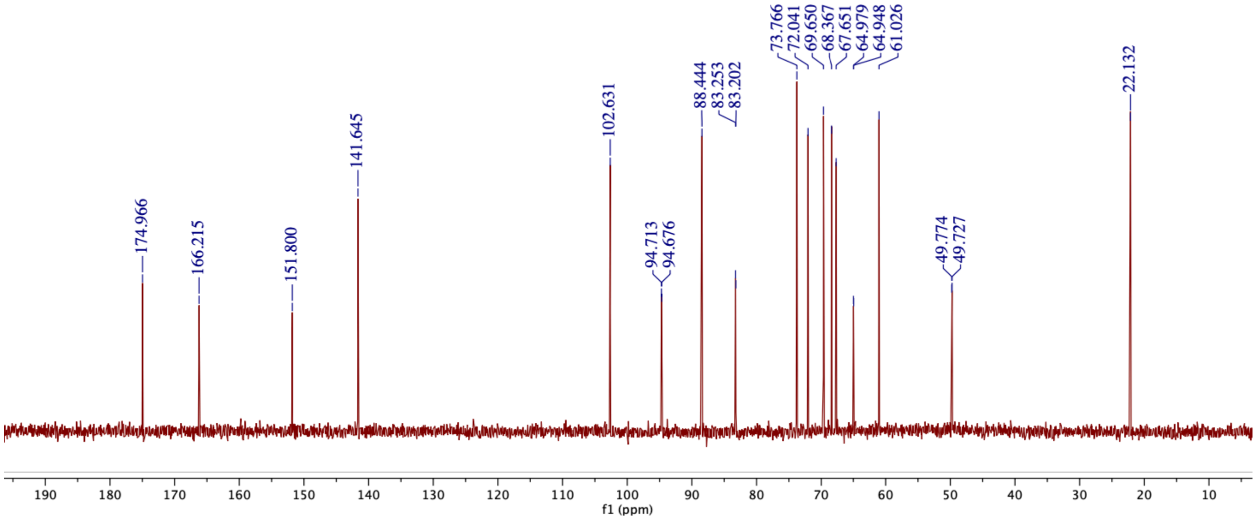

### $^{1}H$ and $^{13}C$ NMR Spectra of UDP-GalNDAz (**8**)

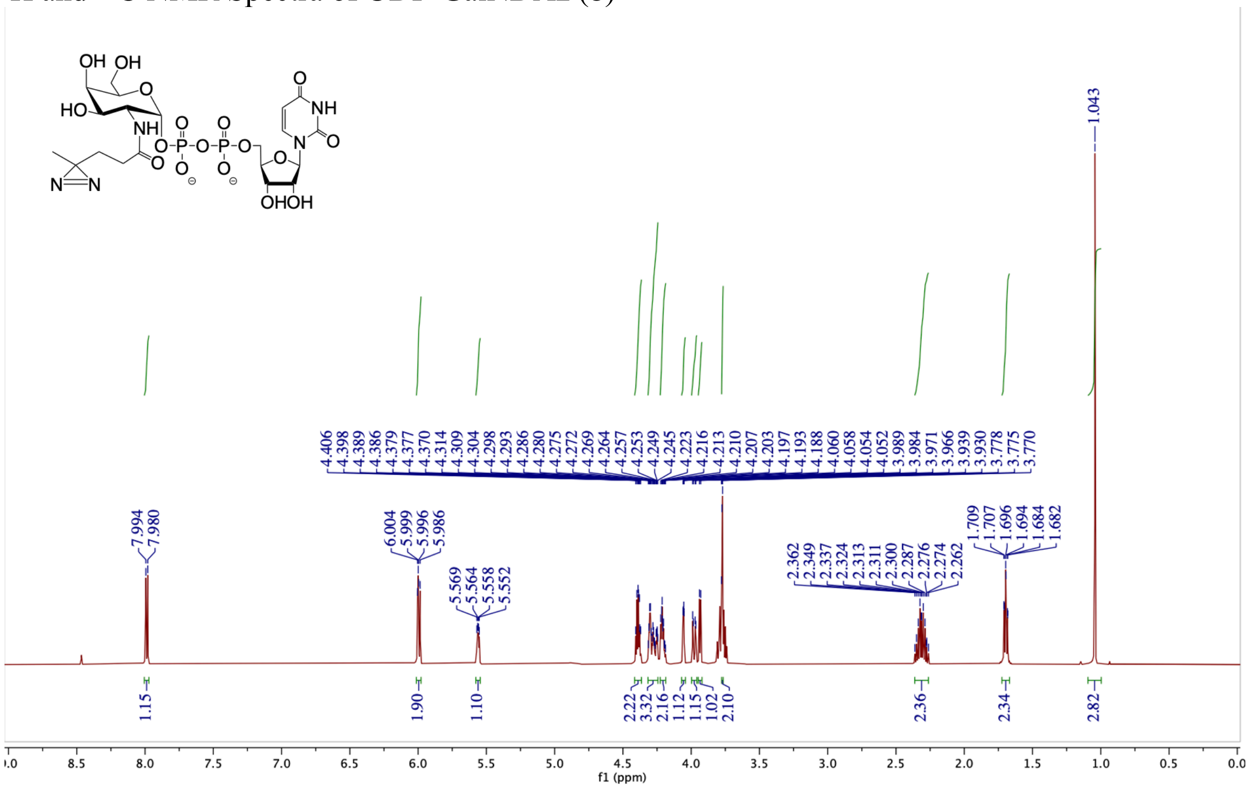

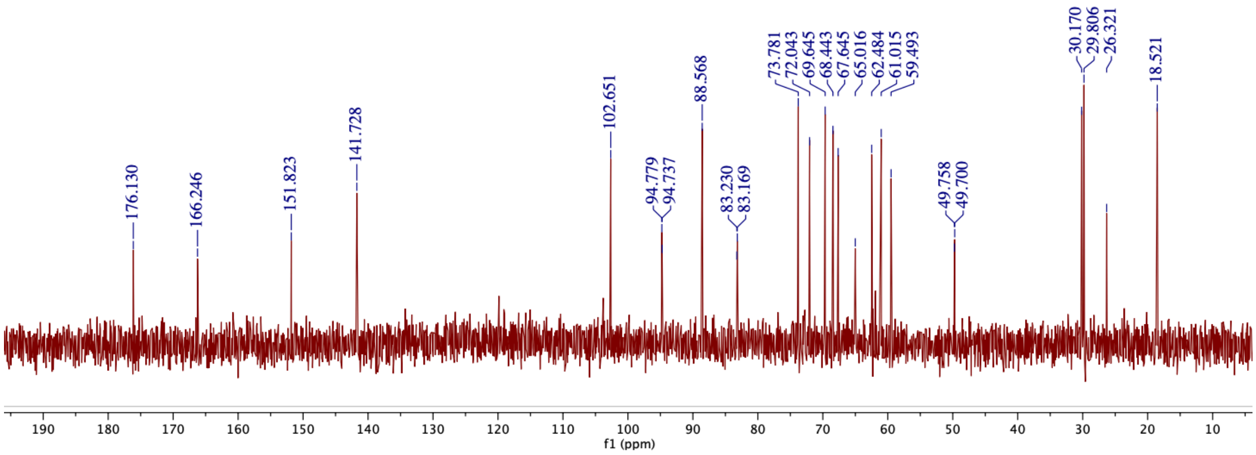
